# A predictive microfluidic model of human glioblastoma to assess trafficking of blood-brain barrier penetrant nanoparticles

**DOI:** 10.1101/2021.12.07.471663

**Authors:** Joelle P. Straehla, Cynthia Hajal, Hannah C. Safford, Giovanni S. Offeddu, Natalie Boehnke, Tamara G. Dacoba, Jeffrey Wyckoff, Roger D. Kamm, Paula T. Hammond

## Abstract

The blood-brain barrier represents a significant challenge for the treatment of high-grade gliomas, and our understanding of drug transport across this critical biointerface remains limited. To advance preclinical therapeutic development for gliomas, there is an urgent need for predictive *in vitro* models with realistic blood-brain barrier vasculature. Here, we report a vascularized human glioblastoma (GBM) model in a microfluidic device that accurately recapitulates brain tumor vasculature with self-assembled endothelial cells, astrocytes, and pericytes to investigate the transport of targeted nanotherapeutics across the blood-brain barrier and into GBM cells. Using modular layer-by-layer assembly, we functionalized the surface of nanoparticles with GBM-targeting motifs to improve trafficking to tumors. We directly compared nanoparticle transport in our *in vitro* platform with transport across mouse brain capillaries using intravital imaging, validating the ability of the platform to model *in vivo* blood-brain barrier transport. We investigated the therapeutic potential of functionalized nanoparticles by encapsulating cisplatin and showed improved efficacy of these GBM-targeted nanoparticles both *in vitro* and in an *in vivo* orthotopic xenograft model. Our vascularized GBM model represents a significant biomaterials advance, enabling in-depth investigation of brain tumor vasculature and accelerating the development of targeted nanotherapeutics.

**Significance Statement:** The blood-brain barrier represents a major therapeutic challenge for the treatment of glioblastoma, and there is an unmet need for *in vitro* models that recapitulate human biology and are predictive of *in vivo* response. Here we present a new microfluidic model of vascularized glioblastoma featuring a tumor spheroid in direct contact with self-assembled vascular networks comprised of human endothelial cells, astrocytes, and pericytes. This model was designed to accelerate the development of targeted nanotherapeutics, and enabled rigorous assessment of a panel of surface-functionalized nanoparticles designed to exploit a receptor overexpressed in tumor-associated vasculature. Trafficking and efficacy data in the *in vitro* model compared favorably to parallel *in vivo* data, highlighting the utility of the vascularized glioblastoma model for therapeutic development.

## Introduction

High-grade gliomas are the most common primary malignant brain tumors in adults (1). These include grade IV astrocytomas, commonly known as glioblastoma multiforme (GBM), which account for more than 50% of all primary brain cancers and have dismal prognoses with a five-year survival rate of less than 5% (2). Due to their infiltrative growth into the healthy brain tissue, surgery often fails to eradicate all tumor cells (3). While chemotherapy and radiation modestly improve median survival (4), most patients ultimately succumb to their tumors. This is primarily due to the presence of a highly selective and regulated endothelium between blood and brain parenchyma known as the blood-brain barrier (BBB) (5), which limits the entry of therapeutics into the brain tissue where tumors are located. The BBB, characterized by a unique cellular architecture of endothelial cells (ECs), pericytes (PCs), and astrocytes (ACs) (6, 7), displays upregulated expression of junctional proteins and reduced paracellular and transcellular transports compared to other endothelia (8). While this barrier protects the brain from toxins and pathogens, it also severely restricts the transport of many therapeutics, as evidenced by the low cerebrospinal fluid-to-plasma ratio of most chemotherapeutic agents (9). There is thus an important need to develop new delivery strategies to cross the BBB and target tumors, enabling sufficient drug exposure (10).

Despite rigorous research efforts to develop effective therapies for high-grade gliomas, the majority of trialed therapeutics have failed to improve outcomes in the clinic even though the agents in question are effective against tumor cells in preclinical models (11). This highlights the inability of current preclinical models to accurately predict the performance of therapeutics in human patients. To address these limitations, we developed an *in vitro* microfluidic model of vascularized GBM tumors embedded in a realistic human BBB vasculature. This BBB-GBM platform features brain microvascular networks (MVNs) in close contact with a GBM spheroid, recapitulating the infiltrative properties of gliomas observed in the clinic (12) and those of the brain tumor vasculature, with low permeability, small vessel diameter, and increased expression of relevant junctional and receptor proteins (7). This platform is well suited for quantifying vascular permeability of therapeutics and simultaneously investigating modes of transport across the BBB and into GBM tumor cells.

There is strong rationale for developing therapeutic nanoparticles (NPs) for GBM and other brain tumors, as they can be used to deliver a diverse range of therapeutic agents, and with appropriate functionalization, can be designed to exploit active transport mechanisms across the BBB (13, 14). Liposomal nanoparticles have been employed in the oncology clinic to improve drug half-life and decrease systemic toxicity (15), but to date, no nanomedicines have been approved for therapeutic indications in brain tumors. We hypothesize that a realistic BBB-GBM model composed entirely of human cells can accelerate preclinical development of therapeutic NPs. Using our BBB-GBM model, we investigated the trafficking of layer-by-layer nanoparticles (LbL-NPs) and ultimately designed a new GBM-targeted NP. The LbL approach leverages electrostatic assembly to generate modular NP libraries with highly controlled architecture. We have used LbL-NPs to deliver a range of therapeutic cargos in pre-clinical tumor models (16, 17) and have recently demonstrated that liposomes functionalized with BBB penetrating ligands improved drug delivery across the BBB to GBM tumors (18). Consistent with clinical data (19), we observed that the low density lipoprotein receptor-related protein 1 (LRP1) was up-regulated in the vasculature near GBM spheroids in the BBB-GBM model and leveraged this information to design and iteratively test a library of NPs. We show that the incorporation of angiopep-2 (AP2) peptide moieties on the surface of LbL-NPs leads to increased BBB permeability near GBM tumors through LRP1-mediated transcytosis. With intravital imaging, we compared the vascular permeabilities of dextran and LbL-NPs in the BBB-GBM platform to those in mouse brain capillaries and validated the predictive potential of our *in vitro* model. Finally, we show the capability of the BBB-GBM platform to screen therapeutic NPs and predict *in vivo* efficacy, demonstrating improved efficacy of cisplatin when encapsulated in GBM-targeting LbL-NPs both *in vitro* and *in vivo*.

## Results

### A vascularized glioblastoma model for the quantification of vascular permeability

To recapitulate the GBM microenvironment and evaluate the transport of therapeutics across the BBB, we developed an *in vitro* BBB-GBM model in a microfluidic device. This platform features a tumor spheroid (GBM spheroid) composed of cells from a patient-derived xenograft (PDX) glioblastoma cell line co-cultured with pericytes, that is embedded in a BBB vascular system in which induced pluripotent stem cell-derived endothelial cells (iPS-ECs), pericytes (PCs), and astrocytes (ACs) self-assemble into perfusable vascular networks (20) (Fig. 1A). We chose to use the GBM22 cell line from the Mayo Clinic Brain Tumor PDX National Resource (21) for these initial studies because it is well-characterized and has been used extensively in preclinical studies as an orthotopic xenograft (22, 23). GBM spheroids grew in close contact with their surrounding vasculature, resulting in a vascularized GBM model after 7 days of culture. The spheroids grew rapidly in the microfluidic devices and were found to co-opt the surrounding BBB vasculature, similar to what is observed in high-grade glioma patients and in animal models of GBM (24, 25) (Fig. S1). Vascular co-option, a process in which tumor cells hijack existing blood vessels, was validated by the similar vessel density measurements in both proximal and distal regions to the GBM spheroid (26) (Fig. S1).

**Figure 1.**
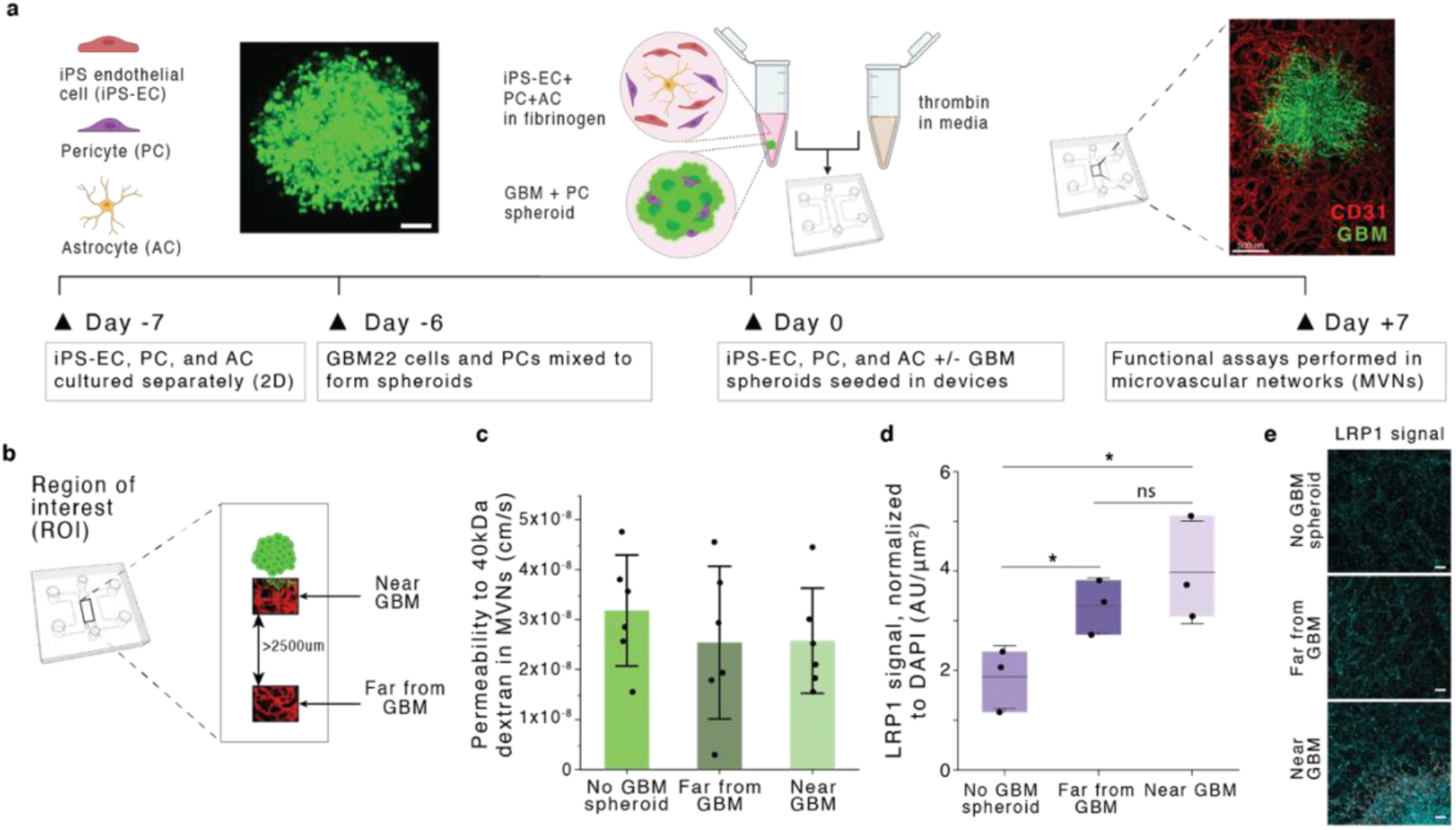
Generation and characterization of a glioblastoma blood-brain barrier microvascular network model (BBB-GBM model). (**A**) Schematic of BBB-GBM formation in a microfluidic device, scale bar=100 μm (left image) and 500 μm (right image). (**B**) Regions of interest (ROIs) identified spatially within the BBB-GBM model, with far from GBM ROIs identified to be at least 2500 μm away from the GBM spheroid. (**C**) Permeability to 40kDa dextran in the vascular networks across different ROI locations, each point represents n=1 device. (**D**) Expression of low density lipoprotein receptor-related protein 1 (LRP1) across different ROI locations, as assessed via immunofluorescence staining, each point represents n=1 device. (**E**) Representative micrographs of LRP1 staining quantified in (D); scale bars=100 μm. In all graphs, bars represent mean ± standard deviation (SD), n.s. denotes not significant, and * p < 0.05. Statistical analyses are described in the Methods section.

Despite a commonly held belief that the BBB is disrupted in human glioblastoma, analyses from patient samples have shown that vessels near high-grade gliomas often exhibit the same integrity and density as vessels found in the healthy brain tissue, depending on their level of infiltration and location within the tumor (27, 28). Particularly, it has been shown that for developing or residual glioma tumors with smaller sizes, the BBB remains intact, and the tumor mass is sustained by normal brain vessels (29). To evaluate our platform in this context, we assessed paracellular permeability to dextran by measuring changes in intensity following fluorescent dextran injection in the vasculature via confocal microscopy. Three regions of interest (ROI) were compared: (i) vessels in devices without GBM tumor (no GBM spheroid), (ii) vessels far away (> 2,500 μm) from the GBM tumor (far from GBM), and (iii) vessels in close proximity to the GBM tumor where co-option is observed (near GBM) (Fig. 1B). We measured no differences in vascular permeability in these three regions, suggesting that tight junctions remain intact even in locations where GBM co-option is evident (Fig. 1C). These findings are striking in comparison with analogous measurements performed by our group using vascularized ovarian or lung tumor models, where vessel permeability was found to be 3-fold larger near the tumor (26). The undisrupted BBB near GBM tumors in this platform attests to the ability of GBM cells to invade and co-opt the vasculature without modifying its properties. This is also demonstrated by unaltered tight and adherens junction protein expressions at endothelial borders in regions of vascular co-option when compared to healthy BBB vessels without GBM tumors (Fig. S1). These results indicate that our BBB-GBM platform is realistic in modeling glioblastoma vasculature, particularly in early tumor development or in recurrent tumor progression following resection where developing tumors are sustained by normal BBB vessels.

The unaltered paracellular permeability of BBB vessels near developing GBM tumors suggests that enhanced localized transport across the BBB via disrupted endothelial junctions is unlikely. Delivery of therapeutics through ligand-based transcellular transport thus offers an avenue for targeted trafficking of therapeutics near GBMs. LRP1, a transport receptor involved in various cellular processes at the BBB, including lipid and lipoprotein metabolism and protease degradation (30), has been shown to be upregulated in GBMs and their surrounding vasculature (31). As a result, there is interest in the design of therapeutics employing LRP1-mediated transport to cross the BBB and specifically target GBM (32–34). We investigated LRP1 expression in the BBB-GBM model and found that the presence of GBM spheroids increases LRP1 expression in vessels both near and far from the spheroid, compared to control devices without tumor (Fig. 1D-E, S1). LRP1 expression was also evidenced within GBM tumor cells. These results guide our subsequent nanoparticle design for enhanced targeted delivery across the BBB near GBM tumors.

### Targeted nanoparticles cross BBB vessels near GBM tumors via LRP1-mediated transport

We next employed the layer-by-layer (LbL) method to develop a NP with enhanced trafficking to GBM cells through tumor-associated vasculature. LbL-NPs consist of a charged nanoparticle core and polyelectrolyte multilayer shell; for this study we started with a liposomal NP core and layered with poly-(L-arginine) and propargyl-modified poly-(L-aspartic acid) (pPLD) as previously described to generate a click-compatible LbL-NP (35). We chose to use a propargyl modification extent of 12% in order to preserve the inherent “stealth” benefits from the anionic, hydrated PLD coating (36) while also incorporating sufficient amounts of targeting ligand. The surface of the NP was functionalized with angiopep-2 (AP2), a peptide designed to target the BBB via interaction with the LRP1 receptor over-expressed in GBM vessels (37) (Fig. 1D-E), generating NPs with favorable size and surface potential for drug delivery applications (Table S1).

We first investigated the NP-cell association of fluorescently-labeled (Cyanine5) bare (Bare NPs), and LbL NPs with an outer surface of pPLD (pPLD NPs) or pPLD functionalized with AP2 (AP2 NPs) (Fig. 2A) with the four cell types in the BBB-GBM model (iPS-ECs, PCs, ACs, and GBM22) using flow cytometry. AP2 NPs exhibited the highest NP-associated fluorescence in all cell lines compared to bare or pPLD NPs and NP internalization in GBM cells was confirmed by microscopy (Fig. S2). The AP2 NP trend was amplified in iPS-ECs, which have high native LRP1 expression and are the first cell type encountered by NPs when crossing the BBB model. We next investigated the ability of NPs to associate with GBM spheroids in the absence of vascular networks to ensure NPs can deliver encapsulated cargo to the cell of interest. Incubation with the different NP formulations showed increased accumulation of AP2 NPs in GBM spheroids compared to bare NPs (Fig. 2B-C).

**Figure 2.**
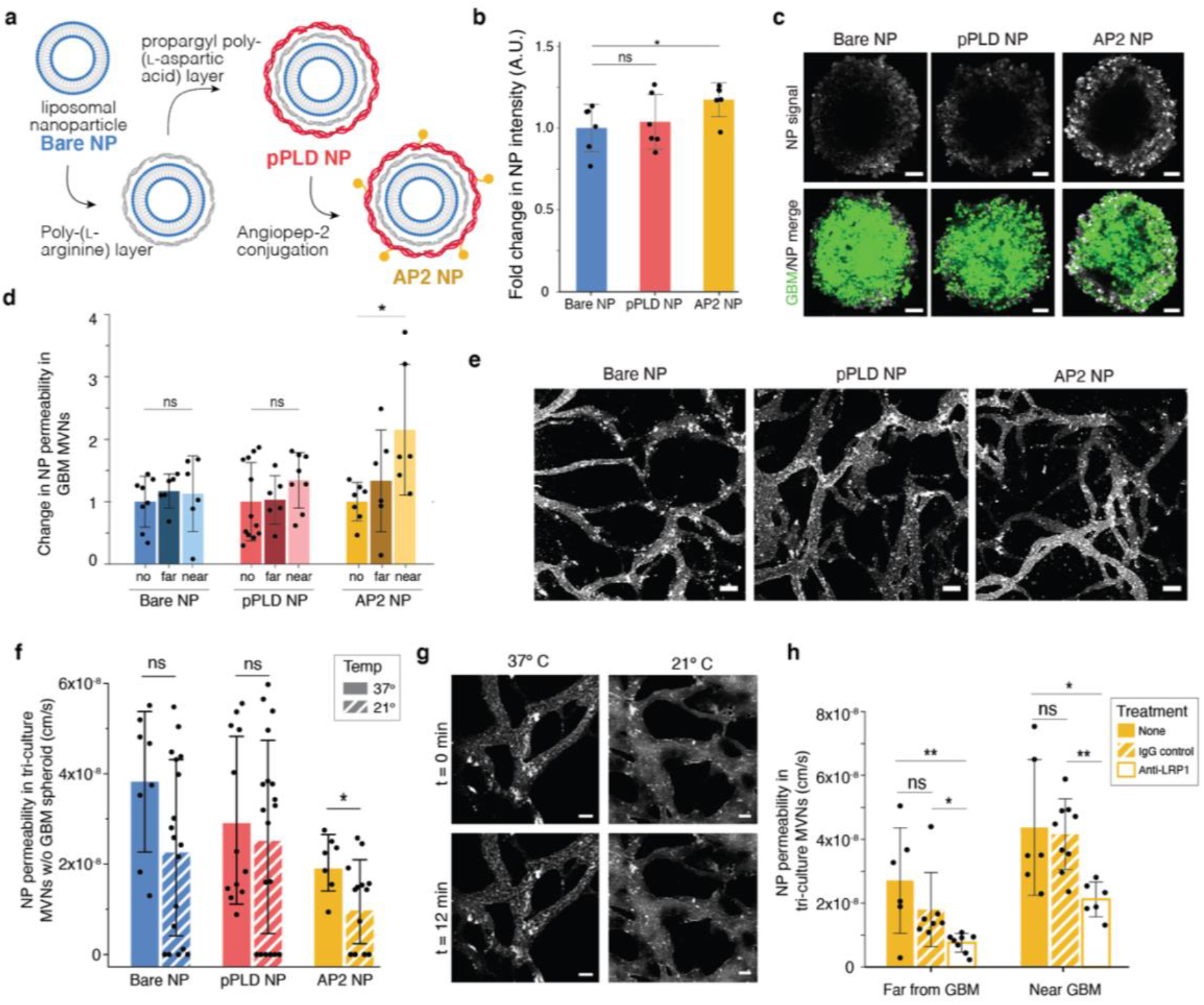
Functionalized LbL-NPs cross BBB microvascular networks near GBM spheroids via LRP1-mediated transport. (**A**) Layer-by-layer assembly of angiopep-2 nanoparticles (AP2-NPs). (**B**) Fold change in mean fluorescence intensity (MFI) of NPs in GBM spheroids without vascular networks, normalized to bare NP; points represents n=1 spheroid. (**C**) Representative GBM spheroids after 12 min NP incubation as quantified in (B); scale bars = 100 μm. (**D**) NP permeability in networks with no GBM spheroid (*no*), and in regions *near* and *far* from a GBM spheroid, normalized to the *no spheroid* device; points represent n = 1 ROI; n = 6 devices/condition were considered. (**E**) Representative images of NPs in the BBB microvessels at t = 0 min following NP perfusion; time-lapse images over 12 min were used to determine permeabilities in (D); scale bars =100 μm. (**F**) BBB vessel permeabilities to the 3 NP formulations in networks without GBM spheroids at 37 and 21 °C. Points represent n = 1 ROI; n = 4 devices per condition were considered. (**G**) Representative images of AP2 NPs at t = 0 and t = 12 min as quantified in (F); scale bars are 50 μm. (**H**) BBB vessel permeabilities to AP2 NPs at 37 °C following incubation for 30 min with anti-LRP1 or IgG control antibodies. Points represent n = 1 ROI; for antibody conditions n = 4 devices were considered; n = 6 non-treated devices. Throughout, bars represent mean ± SD; n.s. denotes not significant, * p < 0.05, and ** p < 0.01. Statistical analyses are described in the Methods section.

NP trafficking was assessed by quantifying vascular permeability as previously done for various therapeutic molecules (38, 39) (Fig. 1C). Of the three NP formulations, AP2 NPs exhibited a significant increase in permeability near the GBM tumor compared to BBB vessels without tumors (Fig. 2D-E, S3). This trend was not only observed with liposomal NPs but also held true with polystyrene NP cores (Fig. S3), indicating that these effects are most likely stemming from the LbL surface functionalization with AP2. In control BBB microvessels (without tumors), bare NPs had slightly higher permeability than pPLD or AP2 NPs (Fig. S3), which we hypothesize to result from two factors. First, the addition of peptides in place of charged groups on the surface may hinder nonspecific transport in a setting with low expression of the targeted receptor. In addition, we identified a size threshold for NP transport across the *in vitro* BBB, and layered functionalized NPs are slightly larger than bare liposome NPs (z-average diameter of 106.2nm for bare NPs, 164.8nm for pPLD NPs, 163.5nm for AP2 NPs; Table S1). To further investigate this size threshold, we chose to use commercially available carboxylated polystyrene NP cores because they are highly uniform in size and amenable to LbL assembly. Testing a range of sizes revealed that non-functionalized NPs ≥100 nm in diameter cannot cross the *in vitro* BBB (Fig. S3). However, the same 100nm-diameter polystyrene NPs with negligible permeability showed a significant increase in permeability near the GBM tumor after LbL surface functionalization with AP2 (Fig. S3). Although polystyrene and liposomal NPs of comparable sizes exhibit similar permeability changes following LbL surface functionalization with AP2, the two NP cores have vastly different physiochemical properties and cross the *in vitro* BBB at different orders of magnitude (permeability of polystyrene NPs ∼ 10^−9^ cm/s compared to ∼ 10^−8^ cm/s for liposomal NPs). This finding is consistent with literature associating the high stiffness of polystyrene nanoparticles (Young’s modulus on the order of ∼10^9^ Pa versus ∼10^6^ Pa for liposomes (40, 41)), with decreased cell internalization due to less efficient internalization mechanisms for stiffer nanomaterials (42–44). In addition, liposomal NP cores are more translationally relevant as they can be used to encapsulate a range of therapeutics for the treatment of GBM tumors.

Having demonstrated increased LRP1 expression in tumor-associated vasculature and increased AP2 NP permeability near GBM spheroids, we hypothesized that AP2 NPs cross the BBB via transcytosis (45), and more specifically via LRP1-mediated transport. An active mode of transport was validated in the *in vitro* BBB vessels by reducing temperatures from 37 °C to 21 °C to prevent vesicle detachment from the cell membrane and transitioning across the cytoplasm to transport NPs from the luminal to abluminal side of the vessels (39). Indeed, permeability of AP2 NPs was reduced at 21 °C, in line with our hypothesis that AP2 NPs cross the BBB via receptor-mediated transcytosis (Fig. 2F-G). As expected, *in vitro* BBB permeability to 40 kDa dextran, which is expected to cross the endothelium via paracellular transport, was not affected by temperature changes (Fig. S3), confirming that reduced temperatures do not affect the functional properties of the endothelium and its paracellular permeability. The bare and pPLD NP formulations also exhibited small, yet not significant, decreases in permeability, suggesting that NPs of this size, regardless of their functionalization, cross the BBB at least in part via vesicular transport (46, 47). These observations are intriguing, and consistent with recently reported findings that the majority of NP transport across the tumor-associated endothelium occurs via active processes (48). Permeability of AP2 NPs was decreased following LRP1 neutralization compared to IgG control in regions near and far from the GBM spheroid, validating AP2 NP shuttling via the LRP1 receptor in the BBB-GBM model (Fig. 2H). These results highlight the testing capabilities of the BBB-GBM model, where NP formulations can be assayed with high spatio-temporal resolution, to identify their transport properties into tumors.

### The in vitro BBB model accurately predicts in vivo permeability

To evaluate the ability of the BBB-GBM platform to mimic the more complex *in vivo* environment, we quantified permeability of dextran and functionalized NPs in mouse capillaries via intravital imaging. Following NP and dextran intravenous administration in animals, time-lapse images were acquired through a cranial window to quantify vascular permeability, as performed in the *in vitro* devices (Fig. 3A, S4, Movie S1). Both dextran and NP signals were clearly observed in mouse cortical capillaries with comparable sizes to *in vitro* BBB vessels (Fig. 3B, S4, Table S2). Dextran permeability values obtained with the imaging and analysis techniques described here were approximately one order of magnitude smaller than measurements performed by other groups in mouse or rat brain capillaries (49, 50) (Table S3). Remarkably, values obtained in mouse BBB vessels were highly consistent with those obtained in the BBB microvascular device for both 10 and 40 kDa dextran (Fig. 3C, S4). Similarly, NP permeabilities corresponded closely to values from the *in vitro* BBB model without tumors (Fig. 3D). In addition to having comparable morphological properties (Table S2), the consistent permeability measurements *in vitro* and *in vivo* highlight the ability of the *in vitro* BBB model to recapitulate functional aspects of the *in vivo* BBB. Of note, permeability studies in mouse BBB capillaries (without tumors) were performed at depths less than 150 μm below the dura, where imaging resolution is optimal. Performing comparable measurements in vessels near GBM tumors *in vivo* would require superficial tumor implantation which is technically challenging and less clinically relevant than an orthotopic model in the deeper regions of the brain, as tumors are likely to extravasate and establish outside the confines of the BBB (51).

**Figure 3.**
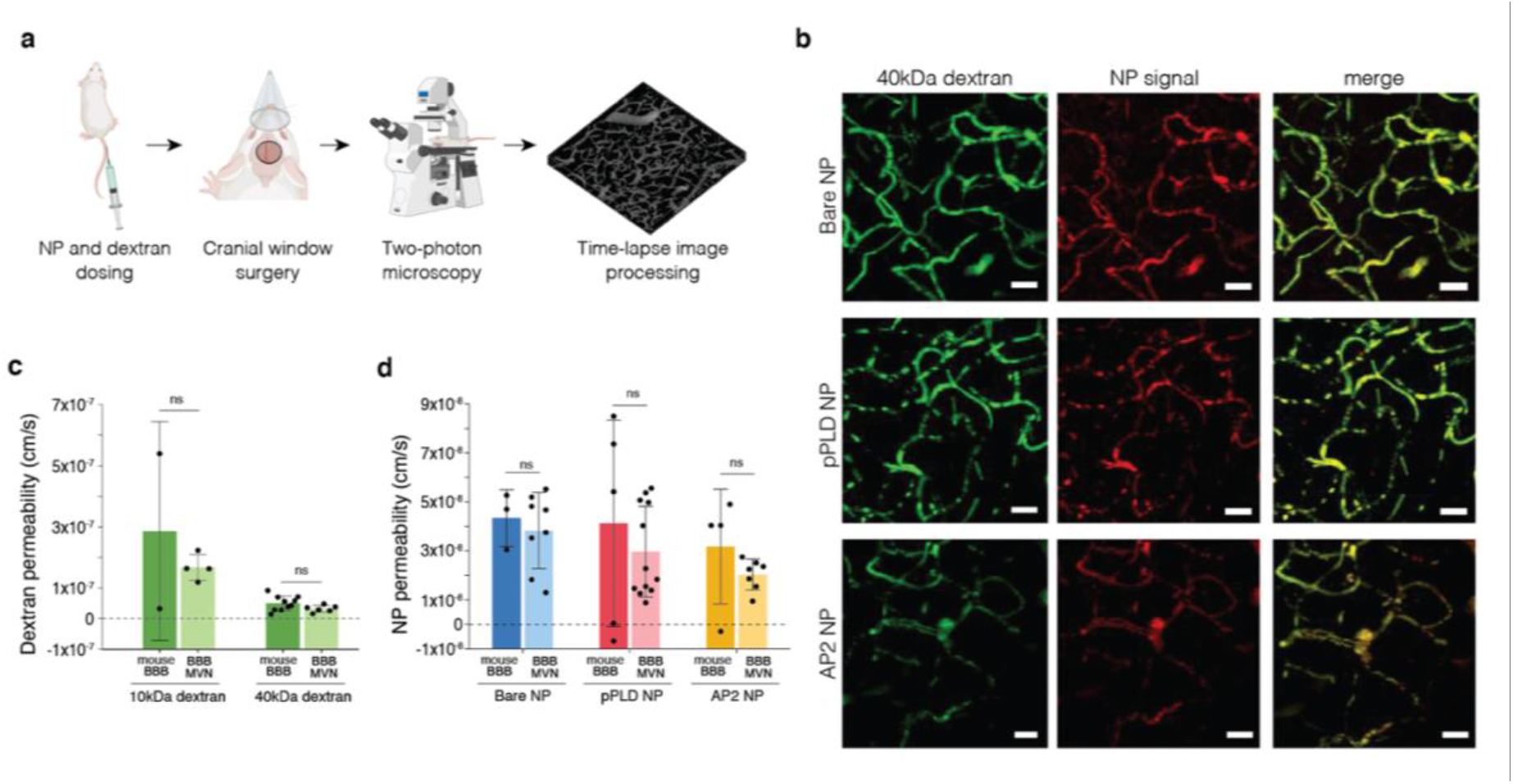
*In vivo* BBB permeability assessed by intravital microscopy is consistent with the *in vitro* BBB model. (**A**) Workflow of intravital imaging in which fluorescent NPs and dextran are dosed systemically and time-lapse imaging is performed in intact brain capillaries. (**B**) Representative images of 40 kDa dextran and NP formulations perfused in mouse BBB capillaries; scale bars = 20 μm. (**C**) BBB permeabilities to fluorescently labeled dextran (10 and 40 kDa) in mouse BBB capillaries and *in vitro* BBB microvessels (no tumors). Points represents n = 1 device; n = 2 mice were considered for 10kDa dextran and n = 10 mice for 40 kDa dextran. (**D**) BBB permeabilities to the 3 NP formulations in mouse BBB capillaries and *in vitro* BBB microvessels (no tumors). Points represent n = 1 ROI; n = 6 independent devices per condition were considered; n = 3-5 mice were considered per condition. Bars represent mean ± SD; n.s. denotes not significant. Statistical analyses are described in the Methods section.

### Therapeutic NPs effectively target tumors in the BBB-GBM model and in vivo

In addition to studying transport, the BBB-GBM model offers the rare opportunity to assess therapeutic efficacy of new agents in the highly relevant setting of microscopic tumor burden. The extent of surgical resection is an important prognostic factor in that achieving a gross total resection portends improved survival for GBM patients, but most tumors recur due to microscopic tumor deposits near the resection cavity (52, 53). To assess the therapeutic potential of LbL-NPs for GBM and the preclinical value of the BBB-GBM model, we next encapsulated the DNA-damaging agent cisplatin (CDDP) in the liposome core of the NPs (Fig. S5). We hypothesized that a selective mode of delivery of CDDP to GBM tumors would lead to improved efficacy and reduced toxicity in the healthy surrounding brain tissue and blood vessels. CDDP was chosen in this study for its nonspecific mechanism of action and its poor BBB penetration (< 0.04 CSF-blood ratio (9)), to evaluate the influence of drug delivery. The BBB-GBM model was instilled with 6 μM of free CDDP, CDDP loaded into bare NPs (bare CDDP NPs, formulated at 7.9 weight% with respect to lipid) or CDDP loaded into AP2 NPs (AP2 CDDP NPs, 4.6 weight%, Table S1). Cisplatin dosing was based on previously determined *in vitro* IC_50_ values for GBM22 cells (Fig. S5) and was continued daily for 4 days via perfusion in BBB microvessels. At the end of the dosing period, all spheroids treated with CDDP containing formulations decreased in size significantly compared to the untreated control, with differences evident as early as 24 hours following initial treatment (Fig. 4A-B).

**Figure 4.**
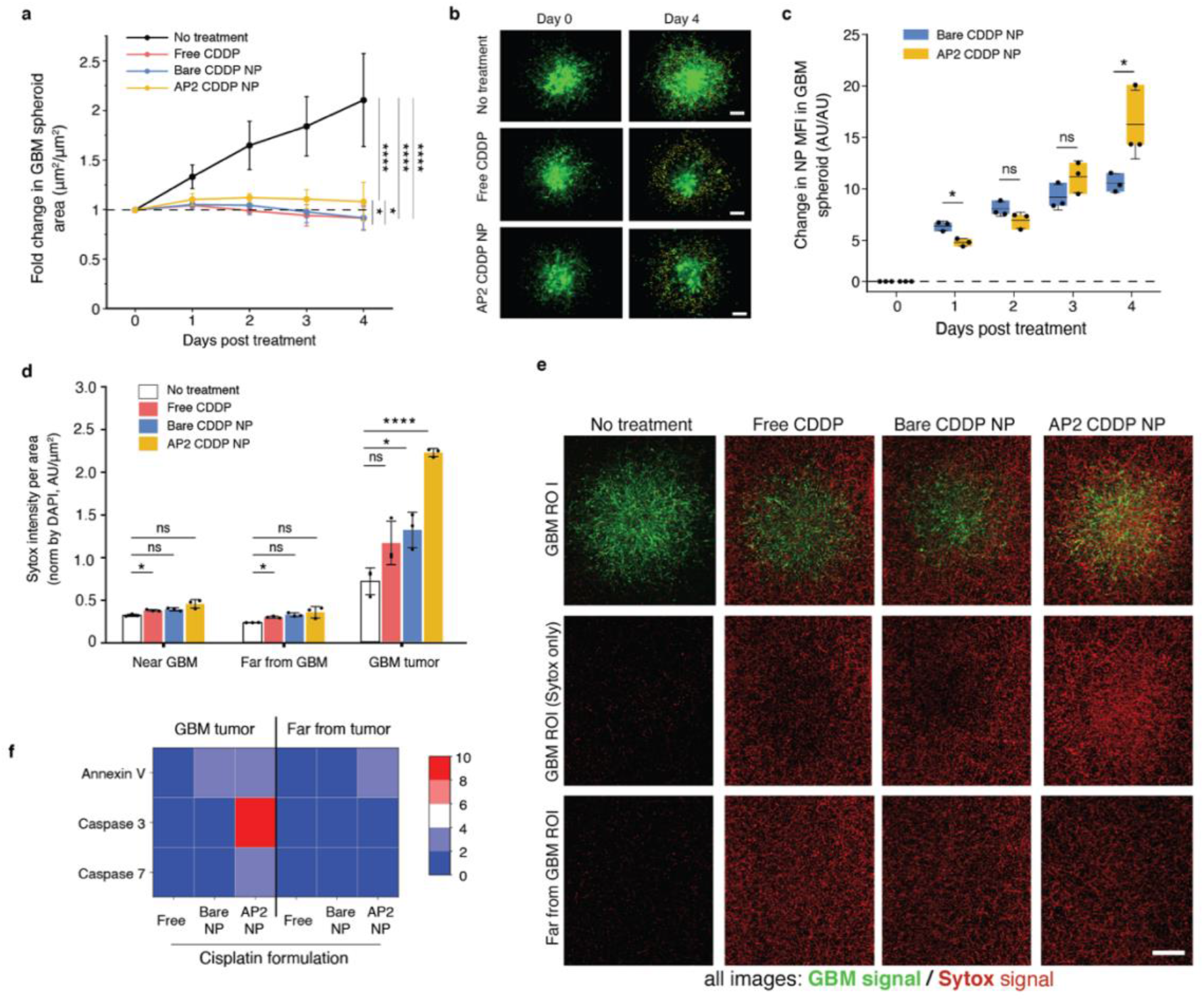
Encapsulation in LbL-NPs improves efficacy and targeted delivery of cisplatin in BBB-GBM. (**A**) GBM spheroid size in BBB-GBM model following treatment with free cisplatin (CDDP), CDDP encapsulated in bare NPs (Bare CDDP NP), or CDDP encapsulated in AP2 NPs (AP2 CDDP NP), compared to untreated devices. Points represent mean ± SD of n=6 devices. (**B**) Representative fluorescent micrographs quantified in (A). Scale bars = 200 μm. (**C**) Change in mean fluorescence intensity (MFI) of NP signal in GBM tumors in the BBB-GBM device over time, following treatment with fluorescently tagged bare-or AP2-CDDP NPs. Points represent n = 1 device. (**D**) MFI of Sytox signal per area (normalized by DAPI) in the 3 ROI locations considered in BBB-GBM devices after treatment with free CDDP, bare CDDP NPs, or AP2 CDDP NPs, and compared to control devices without treatment. Points represent n = 1 device. (**E**) Representative fluorescent micrographs quantified in (D). Scale bars = 500μm (**F**) Heatmap of cell death gene expression levels in 2 ROIs of the devices (inside the GBM tumor and far from the tumor) as quantified by quantitative reverse transcription polymerase chain reaction (qRT-PCR). Bars represent mean ± SD; n.s. denotes not significant, * p < 0.05, ** p < 0.01, *** p < 0.001, and **** p < 0.0001 Statistical analyses are described in the Methods section.

We next assessed trafficking of CDDP-loaded NPs across the BBB and into the tumor space by quantifying NP association with the GBM spheroid over the course of treatment. Bare CDDP NPs initially exhibited higher association with GBM tumors compared to AP2 CDDP NPs; however, this trend was reversed by day 4 of treatment (Fig. 4C). Although all CDDP formulations resulted in tumor growth inhibition, we hypothesized that improved accumulation of AP2 CDDP NPs in GBM cells with repeated dosing may enhance the therapeutic index of the drug and result in reduced cytotoxicity in the local BBB vessels. This was evaluated using the Sytox™ nucleic acid stain to label dead cells in three regions of interest (far, near, and inside GBM spheroids), following treatment with free CDDP, bare CDDP NPs, or AP2 CDDP NPs. Treatment with AP2 CDDP NPs resulted in the largest increase in Sytox signal inside GBM tumors relative to untreated devices (Fig. 4D-E). Near and far from GBM tumors, Sytox signal was minimally increased with all CDDP formulations compared to untreated devices, except for free CDDP which resulted in significant increases in Sytox. Following treatment with the CDDP formulations, regions of the BBB-GBM devices were extracted for quantitative reverse transcription polymerase chain reaction (qRT-PCR). Annexin V, Caspase 3 and Caspase 7 transcripts were all elevated in GBM tumors collected from devices treated with AP2 CDDP NPs compared to the other CDDP formulations (Fig. 4F). There were no major differences in the expression of cell death genes far from the tumor for the different CDDP formulations. Taken together, we show that morphologic size of GBM spheroids is significantly decreased by treatment with CDDP regardless of formulation, but encapsulating CDDP within LbL-NPs with AP2 surface functionalization results in increased and thus more targeted cell death in GBM spheroids without excess damage to the healthy surrounding BBB vasculature, as evidence by comparable Sytox signal far from the tumor across all CDDP formulations.

To test the ability of the BBB-GBM device to predict *in vivo* response, we employed the same CDDP NP formulations in an orthotopic xenograft model generated using the same patient-derived GBM cells used in the BBB-GBM device. To mirror the time frame of the *in vitro* studies, we quantified tumor volume before and after a short dosing period using magnetic resonance imaging (MRI) (Fig. 5A). CDDP formulations were dosed via tail vein every 3 days at a dose of 0.75mg/kg CDDP. This dose was chosen as the highest attainable dose based on limits of passive encapsulation and injection volume, and accounting for conversion from mouse to human (54), is notably lower than the dose employed in GBM clinical trials (55) (2.3mg/m^2^/dose compared to 30mg/m^2^/dose in clinical trial). Despite this limitation, we observed a slower growth trajectory in tumors of animals treated with AP2 CDDP NPs compared to those of animals treated with equivalent CDDP doses in free form (p=0.047) (Fig. 5B), consistent with the improved accumulation of AP2 CDDP NPs in tumors *in vitro*. Levels of cleaved caspase-3 (CC-3) were also increased in tumor tissue after treatment with AP2 CDDP NPs compared to empty liposome control (p=0.0019) (Fig. 5C-D), consistent with increased DNA damage identified in the *in vitro* BBB-GBM device.

**Figure 5.**
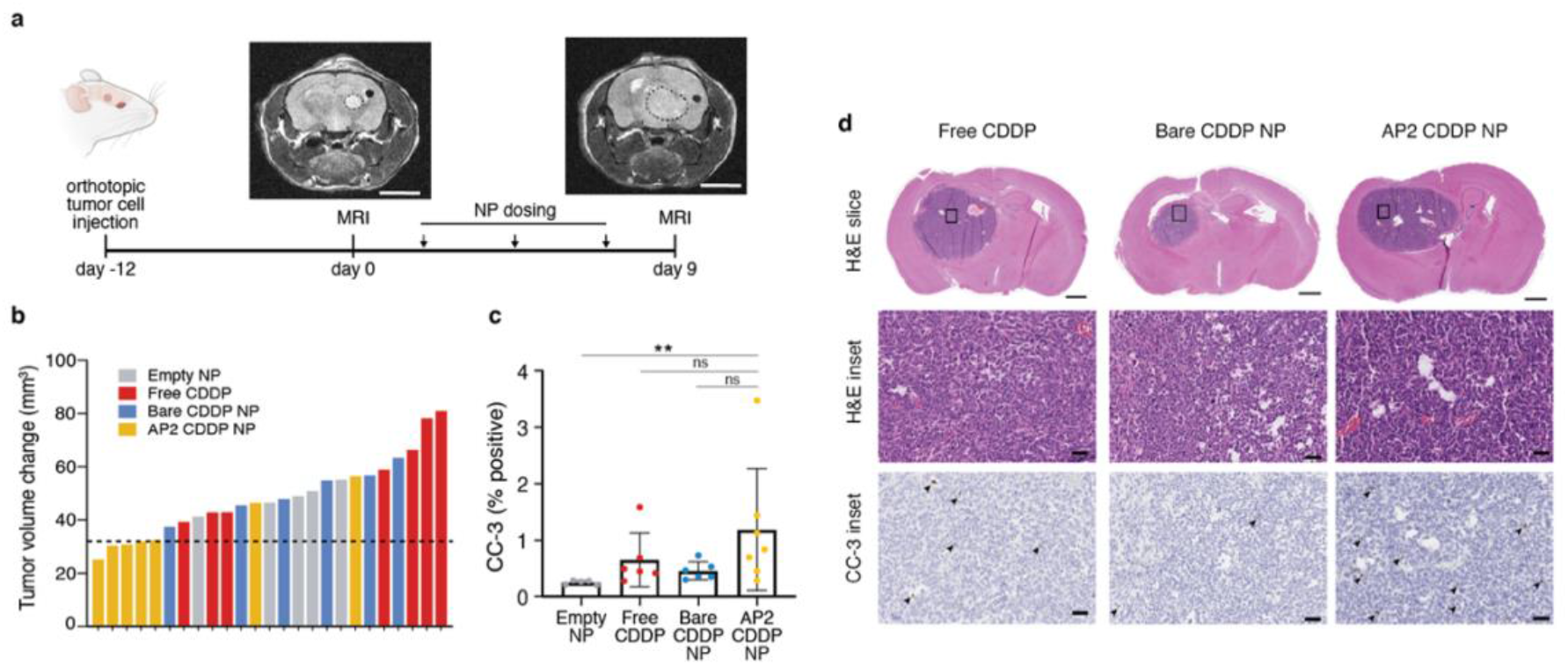
BBB-GBM device predicts differential effects of cisplatin NP formulations in an orthotopic *in vivo* model. (**A**) Timeline of *in vivo* study with orthotopic GBM tumors using magnetic resonance imaging (MRI) to monitor response to therapeutic NPs. Scale bars = 4 mm. (**B**) Waterfall plot for change in tumor volume after treatment on the y-axis, where each bar represents one mouse, dotted line is the median tumor volume change for the AP2 NP group. (**C**) Quantification of cleaved caspase-3 (CC-3) staining in tumor tissue. Each dot represents n = 1 mouse. (**D**) Representative immunohistochemistry micrographs with hematoxylin and eosin (H&E) staining (for context) and CC3 as quantified in c; arrowheads denote CC-3 positive cells. Scale bars = 50 μm except for top row where scale bars = 1 mm. Bars represent mean ± standard deviation (S.D.), n.s. denotes not significant, * p < 0.05, ** p < 0.01. Statistical analyses are described in the Methods section.

Taken together, we show that encapsulation of CDDP in an LbL-NP with AP2 surface functionalization leads to improved efficacy *in vitro* using a BBB-GBM device and *in vivo* using an analogous orthotopic xenograft model. These findings highlight the impact of LbL surface functionalization in nanomedicine and contribute to the existing body of literature supporting AP2 as a promising targeting BBB moiety. Importantly, the *in vitro* human BBB-GBM model allowed us to rigorously interrogate the trafficking and therapeutic effects of multiple NP formulations in a realistic setting predictive of *in vivo* effects. The high spatio-temporal resolution of our *in vitro* BBB-GBM model and its use in dissecting and investigating modes of transport at the BBB make it a valuable pre-clinical testing platform to speed the development of brain tumor-directed therapies.

## Discussion

In this study, we present a new *in vitro* model of the GBM tumor microenvironment that features perfusable human BBB microvessels coming in direct contact with tumor cells. This innovative design provides a robust platform to study the trafficking of tumor-directed therapies across the BBB. With recent advances in *in vitro* technologies and a push for personalized patient models, there has been increased interest in the design of preclinical human GBM assays (56, 57). While existing platforms can recapitulate the many features of GBM tumors, most lack perfusable vasculature (58, 59) or feature tube-like vessels of large diameters that do not come in direct contact with GBM cells (60, 61). Recently, organoid cultures of patient-derived glioma stem cells have been employed to better recapitulate GBM-vascular interactions, however the self-assembled vessels in these cultures are typically not perfusable which limits their use in the context of drug delivery across the BBB and into tumors (62). Here, we address these limitations by incorporating a GBM tumor spheroid into perfusable self-assembled vascular networks composed of iPS-ECs, PCs, and ACs. We have previously demonstrated that these vasculatures recapitulate several properties of the human BBB, including relevant morphology and cellular architecture, low values of permeability, and expression of junction and transport proteins (20, 63). In the BBB-GBM model, spheroids proliferate and infiltrate the surrounding vasculature, resulting in a physiologically relevant model of vascularized GBM tumors co-opting their adjacent vessels as observed in patient brain tumors (64, 24).

Our system has a number of potential applications for the study of the BBB-GBM interface, ranging from fundamental biology to experimental therapeutics. Here, we report one compelling application: the design, optimization, and evaluation of a targeted therapeutic nanoparticle. Leveraging increased expression of LRP1 on blood vessels in proximity to GBM spheroids, we synthesized LbL-NPs with AP2 on the surface and investigated their trafficking across the BBB and into GBM spheroids. Compared to conventional NP design and optimization which generally involves two dimensional assays in one or a few immortalized cell lines (65), we were able to evaluate NP trafficking rapidly and quantitatively in a three-dimensional environment that accurately recapitulates human GBM. We also probed the mechanisms of functionalized NP trafficking with high spatio-temporal resolution and confirmed that our AP2-NPs cross the vasculature via an active process mediated by the LRP1 receptor.

To evaluate the clinical translatability of our *in vitro* model, we performed analogous permeability assays in the murine brain using via intravital imaging and confirmed that our model accurately recapitulates the *in vivo* setting. To our knowledge, direct comparisons of *in vitro* and *in vivo* BBB permeabilities using the same measurement technique have not been previously reported. More importantly, the majority of animal permeability measurements are performed at low resolution (49, 50, 66), which can impact the resulting permeability measurements (39). Our approach circumvented these challenges using three-dimensional volumes containing several interconnected BBB capillaries, resulting in permeability values that matched those of the *in vitro* model with great accuracy. These findings support the continued development of *in vitro* BBB models for translational applications.

In addition to its ability to predict permeability *in vivo*, another important feature of our BBB-GBM platform is its ability to model clinical scenarios with significant treatment challenges, such as residual microscopic tumor deposits after surgery. Even after a gross total resection, highly proliferative and invasive residual tumor cells are often found beyond the margins of resected gliomas, leading to tumor recurrence (67). Given dismal prognosis of recurrent GBM, developing effective treatments for recurrent tumors is of the utmost importance. Our BBB-GBM model, with its barrier properties mimicking the human BBB, provides a new tool to design and evaluate alternative therapeutics targeting residual or recurrent tumors, surrounded by an intact BBB. One of the key advantages of our model is the ability to assess changes in BBB microvasculature –both biologic and functional—in the presence of a patient-derived xenograft tumor spheroid, paving the way for a wide range of basic and translational investigations future work.

We utilized the BBB-GBM model to develop a panel of LbL-NPs and test their trafficking and therapeutic effects, along with untargeted liposomes and polymeric nanoparticles. By functionalizing the surface of cisplatin NPs with AP2, we showed improved NP accumulation and increased apoptosis in the GBM spheroid with minimal damage to surrounding healthy blood vessels, highlighting the potential for rationally designed nanotherapeutics to exert a selective therapeutic effect based on differential trafficking in tumor-associated vasculature. We complemented these studies with analogous investigations in an orthoptic intracranial murine tumor model to determine whether these results correlate with nanoparticle efficacy on a larger scale and observed slower tumor growth in mice treated with AP2 CDDP NPs despite sub-clinical dosing. However, we hypothesize that the rapid growth and large tumor size at the study endpoint mitigated our ability to detect differential therapeutic effects which may be more evident in a microscopic tumor setting. This is a common limitation of *in vivo* GBM studies and is further motivation to develop a predictive *in vitro* model for testing of future, more potent therapies.

With increasing interest in the development of personalized medicine platforms for the testing of therapies, we anticipate that our BBB-GBM model may be able to address this need in the future if paired with patient cells isolated at the time of surgery. Such a model could enable rapid drug screening with both clinical and investigational therapies. Another application of our model may be to address tumor heterogeneity in a systematic manner. Whereas we employed our vascularized GBM model with one PDX tumor model and tested a panel of functionalized nanoparticles, our platform could be similarly employed to compare biologic and functional differences in the vasculature of multiple glioma models.

Our BBB-GBM platform provides a new resource for the scientific community studying GBM and other challenging brain tumors, particularly in the context of drug delivery with targeted NPs. Ultimately, realistic *in vitro* vascularized GBM models can advance our understanding of tumor-blood vessel biology and accelerate the development of brain-penetrant therapeutics.

## Materials and Methods

### Study design

The main objective of this study was to develop a vascularized GBM model and employ it to investigate the trafficking of LbL NPs. All *in vitro* studies utilized at least four devices per group. We validated the results of this study using intravital microscopy in non-tumor bearing mice (n=3-5 per group) and in an orthotopic xenograft mouse model (n=5-7 mice per group).

For all experiments, devices and animals were allocated randomly across different groups. For tumor burden quantification by MRI, all image analyses were performed in a blinded fashion. All animal experiments in this study were approved by the Institutional Animal Care and Use Committee (IACUC) for Massachusetts Institute of Technology and were performed in accordance with the approved guidelines for animal experimentation from the Committee on Animal Care.

### Cell culture and treatments

Human iPS-ECs (Fujifilm Cellular Dynamics, 11713), human brain PCs and ACs (ScienCell), were cultured as described previously (20, 63). The high-grade glioma patient-derived xenograft (PDX) line glioblastoma 22 (GBM22) originated in the Sarkaria lab, Mayo Clinic and cell line identity was confirmed by short tandem repeat (STR) testing.

### Tumor spheroid formation

GBM22 and brain PCs were co-cultured in a low-adhesion 96-well plate (PrimeSurface 96M plate, Sbio) at a ratio of 4:5 to recapitulate tumor-stromal cell ratios commonly observed in solid tumors (68) and ensure the spheroids remain compact in the low-adhesion plates. Spheroids formed over several days by self-aggregation and were cultured for 6-7 days prior to seeding in the microfluidic devices as previously described (20, 63).

### Device fabrication and microvascular network (MVN) formation

The 3-dimensional microfluidic devices employed in this study were fabricated using soft lithography as previously described (64, 69) with dimensions outlined in detail elsewhere (70). Briefly, a larger device with a width of 3 mm for the central gel channel and height of 500 μm was employed to ensure that spheroids can occupy the center of the channel with sufficient space for the formation of surrounding MVNs. To recapitulate the *in vivo* organization of glioma tumors surrounded by brain capillaries, spheroids were carefully removed from the 96-well plate and mixed with iPS-ECs, PCs, and ACs in fibrinogen at the ratios needed to generate the tri-culture BBB MVNs. An equal amount of thrombin was added and mixed with all the cells and spheroid prior to injecting into the devices for fibrin polymerization (69).

### Tumor growth, vessel coverage, and vessel density measurements

Tumor growth in the devices was measured daily between days 0 (seeding) and 7 by quantifying GFP signal of the GBM spheroids using an Eclipse Ti episcope (Nikon) and the Fiji distribution of ImageJ (NIH) (71). Vessel density was computed as previously described (26) using confocal microscopy (FV-1200, Olympus) and staining for CD31 (ab3245, Abcam) at predetermined regions of interest (ROIs): tumor center, proximal, and distal. These results were compared to prior results from our group using human umbilical vein endothelial cell (HUVEC) MVNs and ovarian (Skov3) spheroids or A549 (lung) spheroids (26). Additional details are in the supplemental methods.

### Immunostaining and image analysis

Devices were fixed, permeabilized, blocked prior to staining. Protein visualization was achieved by staining the fixed devices with anti-CD31 (ab3245, Abcam), anti-LRP1 (sc-57351, Santa Cruz Biotechnology), and anti-ZO-1 (61-7300, ThermoFisher Scientific) at 1:200 in PBS, overnight at 4C on a shaker. Secondary antibodies were used at 1:200 in PBS (568 goat anti-rabbit A-11011, or 633 goat anti-mouse A-21052, Invitrogen) and DAPI (D1306, Invitrogen) at 1:1000. Additional staining details are in the supplemental methods. Images were acquired with a confocal laser scanning microscope (FV-1200, Olympus).

For *in vivo* samples, brains were formalin fixed and paraffin embedded before staining with cleaved caspase-3 (CC3 Rabbit Mab, 1:800 #9664L [D175] Cell Signaling Technology) and rabbit polymer secondary (Biocare Medical # RMR 622L). Quantification of CC3 staining was performed in QuPath v0.2.3 (Queen’s University, Belfast, Northern Ireland) using QuPath’s build-in ‘Positive cell detection’ (72) with three regions of interest of the same size manually placed per tumor section.

### Protein expression analysis

Expression analysis of LRP1 was performed using a Proteinsimple automatic western assay as previously described (73) (74) (74)after fixation with PFA. Devices was separated from the glass coverslip using a razor blade, and different regions of the MVNs in fibrin gel were collected: 4 ROIs in the control MVNs without GBM spheroids, and in the MVNs with GBM spheroids, 2 ROIs near and 2 ROIs far from the spheroid.; n=5-6 devices were employed for each condition. Samples were incubated in lysis buffer comprising 10mL of 1X buffer (9803S, Cell Signaling Technologies), 1 μL Benzonase Nuclease (E8263, Millipore Sigma), one tablet of protease inhibitor cocktail (11836170001, Millipore Sigma), and stored at -80 °C. LRP1 signal (sc-57351, Santa Cruz Biotechnology) was normalized to CD31 (ab32457, Abcam) or β-actin (926-42210, Li-Cor) using Compass software v5.0. The output of this automatic western assay is not a standard blot but rather a chemiluminescence spectrum; representative uncropped raw data for LRP1 signal in one device is included in Figure S1.

### Generation and characterization of layer-by-layer nanoparticles with and without cisplatin

Fluorescent liposomes were generated using thin film hydration followed by extrusion as previously described (75), with additional details in the supplemental materials and methods.

For cisplatin-loaded liposomes, the film was rehydrated with a highly concentrated 8mg/ml solution of cisplatin (CDDP) in milliQ water at 80°C prior to extrusion. Next, liposomes were fluorescently labeled through NHS-coupling of sulfo-cyanine5 NHS ester dye (Lumiprobe, Hunt Valley MD) to DSPE headgroups according to the manufacturer instructions. Excess dye and/or drug was removed via KrosFlo II tangential flow filtration (TFF) system (Repligen, Waltham, MA). See supplemental method for cisplatin quantification techniques.

Fluorescent polystyrene cores with carboxylated surfaces were purchased from Invitrogen with diameters of 0.02 μm, 0.1 μm, and 0.5 μm. Layering was achieved by sequentially adding oppositely charged polyelectrolytes as previously described (76) with further details in the supplemental methods. Angiopep-2 was custom synthesized by the Biopolymers and Proteomic Core Facility at MIT with the sequence (K*)TFFYGGSRGKRNNFKTEEY, in which K* denotes modification of the N-terminus with lysine-azide. Copper-based click chemistry was used to conjugate the azide-modified Angiopep-2 to the propargyl-modified PLD as previously described (35), and copper was removed via tangential flow filtration.

The hydrodynamic size, polydispersity index, and surface potential (zeta) were monitored throughout synthesis and prior to downstream experiments using dynamic light scattering and laser doppler anemometry (Malvern ZS90 Particle Analyzer, λ= 633 nm, material/dispersant RI 1.590/1.330; Malvern, Westborough, MA). Negative staining transmission electron microscopy (2100 Field Emission Electron Microscope, JEOL, Peabody MA) with 1% phosphotungstic acid in water was utilized to further characterize cisplatin-loaded liposomes and confirm the dynamic light scattering data.

### NP association with cells in 2-dimensional culture or isolated 3-dimensional spheroids

NP association in cell lines was quantified using flow cytometry (FACS LSR II with HTS sampler, BD Biosciences, San Jose CA) after incubation with NPs at a final concentration of 10μg/ml lipid in media for 24 h. After washing with PBS to remove unassociated NPs, cell suspensions were analyzed for Cy5 fluorescence using a 640nm laser and 670/30 filter. Structured illumination microscopy with image deconvolution was performed after the same NP incubation period and washing steps as for flow cytometry to determine the intracellular localization in GBM22 cells as previously described (77) using an Inverted X71 microscope (Olympus, Center Valley, PA).

NP association in spheroids (without MVNs) was quantified by incubating the spheroids with 80 μL of 30ucg/ml (wrt lipid concentration) NP suspension (bare, pPLD, or AP2 NPs) for 12 minutes. Day 13 was chosen to ensure that the spheroids in 3-dimensions without MVNs were assessed at the same time as spheroids in the MVNs. Following NP incubation, spheroids were washed with PBS and placed on a glass coverslip for 3-dimensional imaging with a confocal microscope (FV-1200, Olympus). NP intensity into the spheroids of ∼ 600 μm in diameter was quantified by averaging the intensity per location of four diametrical lines in the spheroid using the “Plot Profile” function of ImageJ.

### In vitro permeability assay in the BBB-GBM model

To prevent dye leakage from the side channels, a monolayer of iPS-ECs was added to both media channels of the microfluidic device on day 4 following cell seeding (38). Permeability was measured between days 6 and 8 in the MVNs with and without tumor spheroids. The MVN permeabilities to 10 and 40 kDa dextran, polystyrene NPs (bare, pPLD, and AP2), as well as liposome NPs (bare, pPLD, and AP2) were quantified following perfusion of 80 μL suspensions as previously described (38). Bare polystyrene NPs of different sizes were also employed to assess size-dependent transport across the *in vitro* BBB MVNs. Briefly, devices were imaged via confocal microscopy (FV-1200, Olympus) at 12-minute intervals in an environmental chamber maintained at 37C and 5% CO2. Following automatic thresholding and segmentation using the Fiji distribution of ImageJ, z-stack images were employed to generate a 3-dimensional mask of the microvasculature(38) (38). Analysis of NP or dextran transport across the BBB (with and without tumor spheroids) was performed as previously described (38).

### Cisplatin NP treatment in vitro and cell death quantification

Cisplatin (CDDP) NPs were made fresh and cisplatin concentrations in each NP formulation were quantified prior to each experiment. BBB-GBM devices were treated daily with 6 μM free CDDP or CDDP NPs per day. Tumor size was measured over time as described above. Fluorescently-labeled CDDP NPs with cyanine5 were used to quantify CDDP NP uptake in GBM tumors. To evaluate cell death, devices were incubated with 5 μM of Sytox Orange (Catalog # S11368, ThermoFisher Scientific) and DAPI for 1 h, applying a hydrostatic pressure drop across the gel channel (69). Confocal microscope images (FV-1200) were automatically thresholded and segmented using the Fiji distribution of ImageJ, z-stack images were employed to generate a 3-dimensional mask of the GBM tumors to quantify Sytox signal inside and near the tumor(38) (38).

### Real-time quantitative reverse transcription polymerase chain reaction (qRT-PCR)

Gene expression was quantified via real-time qRT-PCR for BBB-GBM devices treated with free CDDP, bare CDDP NPs, or AP2 CDDP NPs using an RNeasy Mini Kit (74104, Qiagen) and the 7900HT Fast Real-Time PCR System using the TaqMan Fast Advanced Master Mix (4444556, Thermo Fisher Scientific) as previously described (63). Additional details are in the supplemental methods.

### Intravital imaging and mouse permeability assay

For intravital imaging, NCR/nude mice (Taconic, Rensselaer, NY) were injected with fluorescent, functionalized liposomal NPs (100μL via tail vein at a concentration of 0.5mg/ml lipid in 5% dextrose), then anesthetized according IACUC-approved protocol. Immediately prior to cranial window surgery, mice were dosed with fluorescent dextran of varying molecular weights (100μL via retro-orbital injection at a concentration of 2mg/ml dextran). To create the cranial window, the skull was exposed and a high-speed hand drill (Dremel) was used to thin the skull until the dura mater was exposed over the right frontal cortex. Multiphoton imaging was performed on an Olympus FV-1000MPE multiphoton microscope (Olympus) using a 25X, N.A. 1.05 objective. Excitation was achieved using a femtosecond pulse laser at 840 nm, and emitted fluorescence was collected by PMTs with emission filters of 425/30 nm for Collagen 1, 525/45 nm for FITC-labeled dextran and 672/30 nm for Cy5 NPs. Collagen 1 was excited by second harmonic generation and emits as polarized light at half the excitation wavelength. The collagen 1 signal was used to identify the dura such that the vessels imaged were within the cortex (50-100μm below the dura, Image S4). Images were acquired every 1-2 minutes for 10-20 minutes for analysis, as described below. Mice were maintained under anesthesia for the duration of the imaging and then humanely euthanized.

Acquired images from intravital imaging were then thresholded and segmented using the Fiji distribution of ImageJ just as in the *in vitro* permeability workflow described above. Vessels below the dura and arteries were considered to ensure that these represent BBB capillaries in the mouse brain. The microvasculature filled with dextran (dextran channel) was employed to generate a 3-dimensional mask of the BBB mouse vessels (see Fig. 3A and Fig. S6). This mask was employed to analyze both dextran and nanoparticle transport since dextran filled the entire vasculature and resulted in the most accurate mask of the 3-dimensional vessels. After masking, analysis of NP or dextran transport was performed as previously described (38).

### Vessel dimensions

Acquired images from *in vitro* and *in vivo* samples perfused with fluorescent dextran were analyzed for vessel dimensions. Z-stack images were thresholded and segmented and the built-in skeletonize function of Fiji was used to measure vessel dimensions as previously described (70).

### Tumor implantation and nanoparticle treatment

For tumor implantation, we utilized a modified version of the intracranial xenograft protocol developed in the lab of Dr. Jann Sarkaria (78). In brief, NCR/nude mice (Taconic, Rensselaer, NY) were anesthetized with ketamine and xylazine and placed in a stereotactic head frame (Stoelting, Wood Dale, IL). Using sterile technique, the skull was exposed and a small burr hold was made at coordinates 1mm lateral and 2mm posterior to Bregma. A total of 200,000 cells were injected 3mm below the dura using a 33G 5ul Neuros syringe (Hamilton Company, Reno, NV) and Stoelting quintessential stereotaxic injector at a rate of 1ul/minute. Twelve days after tumor implantation, mice with confirmed intracranial tumors by MRI were randomized and treated with three doses of CDDP-containing NPs or free drug (0.75mg CDDP/kg/dose). Control mice received empty control nanoparticles with equivalent lipid to CDDP NP. All solutions were suspended in 5% dextrose and dosed via tail vein at 100uL/injection.

### MRI methods and tumor volume quantification

Magnetic Resonance Imaging (MRI) was performed on a former Varian/Agilent 7T MRI operated by Bruker AV4 NeoBioSpec70-20USR console, equipped with a Bruker QSN075/040 RF coil. Mice were anesthetized with isoflurane throughout according to approved IACUC protocol. Data were collected and reconstructed within Bruker Paravision PV360 v2.0. T2 Weighted images were obtained using TurboRARE sequence with the following parameters: axial orientation, TR/TE=3000/25 ms, 256×256 matrix, field of view (FOV)=20×20mm^2^, interleaved number of slices=32, no gap and slice thickness=0.5mm, number of averages=4, RARE factor 8. Images were converted to DICOM format.

MRI images were analyzed using the Fiji distribution of ImageJ using a published method for diameter based measurement (79). Using the native images obtained in the coronal plane, the z-slice with maximum craniocaudal (d_cc_) and lateral (d_l_) dimensions was determined, and these diameters recorded. An axial reconstruction was then generated using the built-in ‘Reslice’ function in Fiji with output spacing of 0.5mm (z-slice distance), creating a 40 slice axial image. Using axial images, the slice with maximal anteroposterior diameter (d_ap_) determined. The diameter-based volume (V) was then computed using the ellipsoid formula (V = d_cc_ × d_l_ × d_ap_ × Π / 6). On MRI imaging, some mice were noted to have small subcutaneous collections consistent with tumor cell extravasation from the burr hole. In these cases, tumors were only considered evaluable if there was a clear and distinct intracranial component and were excluded from all analyses otherwise.

## Statistical analysis

All data are plotted as mean ± standard deviation (SD), unless indicated otherwise. Statistical significance was assessed using student’s t-tests when comparing two conditions/groups, one-way analysis of variance (ANOVA) with Tukey’s honestly significant difference (HSD) post-hoc test when comparing > 2 groups, or Kruskal-Wallis multiple comparison test (when applicable) with the software Prism® (GraphPad). For non-homogeneity of variances as determined via Levene’s test, Brown-Forsythe and Welch ANOVA with Dunnett’s T3 post-hoc test was performed with the software Prism®. Results were represented as follows: n.s. stands for not significant, * denotes p < 0.05, ** denotes p < 0.01, *** denotes p < 0.001, and **** denotes p < 0.0001. In all *in vitro* experiments, 6-8 devices per condition were employed unless otherwise indicated. In all *in vivo* mice experiments, 3-7 mice per condition were used unless otherwise indicated.

## Acknowledgments

We thank the Koch Institute’s Robert A. Swanson (1969) Biotechnology Center (supported by NCI grant P30-CA14051) for technical support, specifically the Microscopy Core Facility, Hope Babette Tang (1983) Histology Core Facility, Peterson (1957) Nanotechnology Materials Core Facility, Flow Cytometry Core Facility, Preclinical Modeling, Imaging & Testing Core Facility, and Biopolymers & Proteomics Core Facility. Figures 1, 3, and 5 created in part with BioRender.com. We also thank all members of the Kamm and Hammond laboratories for helpful discussions while developing this work.

## References

1. Q. T. Ostrom, et al., CBTRUS Statistical Report: Primary Brain and Other Central Nervous System Tumors Diagnosed in the United States in 2012–2016. Neuro-Oncology 21, v1–v100 (2019).

2. Q. T. Ostrom, D. J. Cote, M. Ascha, C. Kruchko, J.S. Barnholtz-Sloan, Adult Glioma Incidence and Survival by Race or Ethnicity in the United States From 2000 to 2014. JAMA Oncol 4, 1254–1262 (2018).

3. M. Glas, et al., Residual tumor cells are unique cellular targets in glioblastoma. Ann Neurol 68, 264–9 (2010).

4. R. Stupp, et al., Radiotherapy plus concomitant and adjuvant temozolomide for glioblastoma. N Engl J Med 352, 987–96 (2005).

5. N. J. Abbott, A. A. Patabendige, D. E. Dolman, S. R. Yusof, D. J. Begley, Structure and function of the blood-brain barrier. Neurobiol Dis 37, 13–25 (2010).

6. N. J. Abbott, L. Rönnbäck, E. Hansson, Astrocyte-endothelial interactions at the blood-brain barrier. Nat Rev Neurosci 7, 41–53 (2006).

7. C. Hajal, M. Campisi, C. Mattu, V. Chiono, R. D. Kamm, Models of molecular and nano-particle transport across the blood-brain barrier. Biomicrofluidics 12, 042213 (2018).

8. D. S. Hersh, et al., Evolving Drug Delivery Strategies to Overcome the Blood Brain Barrier. Curr Pharm Des 22, 1177–1193 (2016).

9. L. L. Muldoon, et al., Chemotherapy delivery issues in central nervous system malignancy: a reality check. J Clin Oncol 25, 2295–2305 (2007).

10. R. K. Oberoi, et al., Strategies to improve delivery of anticancer drugs across the blood-brain barrier to treat glioblastoma. Neuro Oncol 18, 27–36 (2016).

11. M. Touat, A. Idbaih, M. Sanson, K. L. Ligon, Glioblastoma targeted therapy: updated approaches from recent biological insights. Ann Oncol 28, 1457–1472 (2017).

12. C. D. Arvanitis, G. B. Ferraro, R. K. Jain, The blood-brain barrier and blood-tumour barrier in brain tumours and metastases. Nat Rev Cancer 20, 26–41 (1).

13. Y. Zhou, Z. Peng, E. S. Seven, R. M. Leblanc, Crossing the blood-brain barrier with nanoparticles. J Control Release 270, 290–303 (01 28).

14. C. Ferraris, R. Cavalli, P. P. Panciani, L. Battaglia, Overcoming the Blood-Brain Barrier: Successes and Challenges in Developing Nanoparticle-Mediated Drug Delivery Systems for the Treatment of Brain Tumours. Int J Nanomedicine 15, 2999–3022 (2020).

15. D. Bobo, K. J. Robinson, J. Islam, K. J. Thurecht, S. R. Corrie, Nanoparticle-Based Medicines: A Review of FDA-Approved Materials and Clinical Trials to Date. Pharm Res 33, 2373–2387 (2016).

16. E. C. Dreaden, et al., Tumor-Targeted Synergistic Blockade of MAPK and PI3K from a Layer-by-Layer Nanoparticle. Clin Cancer Res 21, 4410–9 (2015).

17. L. Gu, Z. J. Deng, S. Roy, P. T. Hammond, A Combination RNAi-Chemotherapy Layer-by-Layer Nanoparticle for Systemic Targeting of KRAS/P53 with Cisplatin to Treat Non-Small Cell Lung Cancer. Clin Cancer Res 23, 7312–7323 (2017).

18. F. C. Lam, et al., Enhanced efficacy of combined temozolomide and bromodomain inhibitor therapy for gliomas using targeted nanoparticles. Nat Commun 9, 1991 (5).

19. K. Boyé, et al., The role of CXCR3/LRP1 cross-talk in the invasion of primary brain tumors. Nat Commun 8, 1571 (11).

20. M. Campisi, et al., 3D self-organized microvascular model of the human blood-brain barrier with endothelial cells, pericytes and astrocytes. Biomaterials 180, 117–129 (10).

21. R. A. Vaubel, et al., Genomic and Phenotypic Characterization of a Broad Panel of Patient-Derived Xenografts Reflects the Diversity of Glioblastoma. Clin Cancer Res 26, 1094–1104 (3).

22. J. L. Pokorny, et al., The Efficacy of the Wee1 Inhibitor MK-1775 Combined with Temozolomide Is Limited by Heterogeneous Distribution across the Blood-Brain Barrier in Glioblastoma. Clin Cancer Res 21, 1916–24 (2015).

23. G. J. Kitange, et al., Inhibition of histone deacetylation potentiates the evolution of acquired temozolomide resistance linked to MGMT upregulation in glioblastoma xenografts. Clin Cancer Res 18, 4070–9 (2012).

24. G. J. Baker, et al., Mechanisms of glioma formation: iterative perivascular glioma growth and invasion leads to tumor progression, VEGF-independent vascularization, and resistance to antiangiogenic therapy. Neoplasia 16, 543–61 (2014).

25. E. A. Kuczynski, P. B. Vermeulen, F. Pezzella, R. S. Kerbel, A. R. Reynolds, Vessel co-option in cancer. Nat Rev Clin Oncol 16, 469–493 (8).

26. K. Haase, G. S. Offeddu, M. R. Gillrie, R. D. Kamm, Endothelial Regulation of Drug Transport in a 3D Vascularized Tumor Model. Adv Funct Mater 30 (2020).

27. J. N. Sarkaria, et al., Is the blood-brain barrier really disrupted in all glioblastomas? A critical assessment of existing clinical data. Neuro Oncol 20, 184–191 (1).

28. E. K. Nduom, C. Yang, M. J. Merrill, Z. Zhuang, R. R. Lonser, Characterization of the blood-brain barrier of metastatic and primary malignant neoplasms. J Neurosurg 119, 427–33 (2013).

29. L. G. Dubois, et al., Gliomas and the vascular fragility of the blood brain barrier. Front Cell Neurosci 8, 418 (2014).

30. S. L. Gonias, W. M. Campana, LDL receptor-related protein-1: a regulator of inflammation in atherosclerosis, cancer, and injury to the nervous system. Am J Pathol 184, 18–27 (2014).

31. M. Demeule, et al., Involvement of the low-density lipoprotein receptor-related protein in the transcytosis of the brain delivery vector angiopep-2. J Neurochem 106, 1534–44 (2008).

32. C. Ché, et al., New Angiopep-modified doxorubicin (ANG1007) and etoposide (ANG1009) chemotherapeutics with increased brain penetration. J Med Chem 53, 2814–24 (2010).

33. Y. Bertrand, et al., Influence of glioma tumour microenvironment on the transport of ANG1005 via low-density lipoprotein receptor-related protein 1. Br J Cancer 105, 1697–707 (2011).

34. X. Ji, H. Wang, Y. Chen, J. Zhou, Y. Liu, Recombinant expressing angiopep-2 fused anti-VEGF single chain Fab (scFab) could cross blood-brain barrier and target glioma. AMB Express 9, 165 (2019).

35. N. Boehnke, et al., Theranostic Layer-by-Layer Nanoparticles for Simultaneous Tumor Detection and Gene Silencing. Angew Chem Int Ed Engl 59, 2776–2783 (2020).

36. S. Correa, E. C. Dreaden, L. Gu, P. T. Hammond, Engineering nanolayered particles for modular drug delivery. J Control Release 240, 364–386 (2016).

37. M. Demeule, et al., Identification and design of peptides as a new drug delivery system for the brain. J Pharmacol Exp Ther 324, 1064–72 (2008).

38. G. S. Offeddu, et al., An on-chip model of protein paracellular and transcellular permeability in the microcirculation. Biomaterials 212, 115–125 (2019).

39. G. S. Offeddu, et al., Application of Transmural Flow Across In Vitro Microvasculature Enables Direct Sampling of Interstitial Therapeutic Molecule Distribution. Small 15, e1902393 (2019).

40. X. Liang, G. Mao, K. Y. Ng, Mechanical properties and stability measurement of cholesterol-containing liposome on mica by atomic force microscopy. J Colloid Interface Sci 278, 53–62 (2004).

41. D. Guo, J. Li, G. Xie, Y. Wang, J. Luo, Elastic Properties of Polystyrene Nanospheres Evaluated with Atomic Force Microscopy: Size Effect and Error Analysis. Langmuir 30, 7206–7212 (2014).

42. R. Hartmann, M. Weidenbach, M. Neubauer, A. Fery, W. J. Parak, Stiffness-dependent in vitro uptake and lysosomal acidification of colloidal particles. Angew Chem Int Ed Engl 54, 1365–8 (2015).

43. L. Ribovski, et al., Low nanogel stiffness favors nanogel transcytosis across an in vitro blood– brain barrier. Nanomedicine: Nanotechnology, Biology and Medicine 34, 102377 (2021).

44. P. Guo, et al., Nanoparticle elasticity directs tumor uptake. Nat Commun 9, 130 (01 09).

45. Y. Bertrand, et al., Transport characteristics of a novel peptide platform for CNS therapeutics. J Cell Mol Med 14, 2827–39 (2010).

46. I. Canton, G. Battaglia, Endocytosis at the nanoscale. Chem Soc Rev 41, 2718–39 (2012).

47. B. Oller-Salvia, M. Sánchez-Navarro, E. Giralt, M. Teixidó, Blood-brain barrier shuttle peptides: an emerging paradigm for brain delivery. Chem Soc Rev 45, 4690–707 (2016).

48. S. Sindhwani, et al., The entry of nanoparticles into solid tumours. Nat Mater 19, 566–575 (5).

49. W. Yuan, Y. Lv, M. Zeng, B. M. Fu, Non-invasive measurement of solute permeability in cerebral microvessels of the rat. Microvasc Res 77, 166–73 (2009).

50. N. Kutuzov, H. Flyvbjerg, M. Lauritzen, Contributions of the glycocalyx, endothelium, and extravascular compartment to the blood-brain barrier. Proc Natl Acad Sci U S A 115, E9429–E9438 (10).

51. M. Patrizii, M. Bartucci, S. R. Pine, H. E. Sabaawy, Utility of Glioblastoma Patient-Derived Orthotopic Xenografts in Drug Discovery and Personalized Therapy. Front Oncol 8, 23 (2018).

52. T. J. Brown, et al., Association of the Extent of Resection With Survival in Glioblastoma: A Systematic Review and Meta-analysis. JAMA Oncol 2, 1460–1469 (2016).

53. M. Rapp, et al., Recurrence Pattern Analysis of Primary Glioblastoma. World Neurosurg 103, 733–740 (2017).

54. A. B. Nair, S. Jacob, A simple practice guide for dose conversion between animals and human. J Basic Clin Pharm 7, 27–31 (2016).

55. J. C. Buckner, et al., Phase III trial of carmustine and cisplatin compared with carmustine alone and standard radiation therapy or accelerated radiation therapy in patients with glioblastoma multiforme: North Central Cancer Treatment Group 93-72-52 and Southwest Oncology Group 9503 Trials. J Clin Oncol 24, 3871–9 (2006).

56. S. Pozzi, et al., Meet me halfway: Are in vitro 3D cancer models on the way to replace in vivo models for nanomedicine development? Adv Drug Deliv Rev 175, 113760 (2021).

57. D. Sood, et al., 3D extracellular matrix microenvironment in bioengineered tissue models of primary pediatric and adult brain tumors. Nat Commun 10, 4529 (10).

58. W. Diao, et al., Behaviors of Glioblastoma Cells in in Vitro Microenvironments. Sci Rep 9, 85 (1).

59. M. T. Ngo, B. A. Harley, The Influence of Hyaluronic Acid and Glioblastoma Cell Coculture on the Formation of Endothelial Cell Networks in Gelatin Hydrogels. Adv Healthc Mater 6 (2017).

60. H. Xu, et al., A dynamic in vivo-like organotypic blood-brain barrier model to probe metastatic brain tumors. Sci Rep 6, 36670 (11).

61. M. S. Ozturk, et al., High-resolution tomographic analysis of in vitro 3D glioblastoma tumor model under long-term drug treatment. Sci Adv 6, eaay7513 (3).

62. A. Linkous, et al., Modeling Patient-Derived Glioblastoma with Cerebral Organoids. Cell Reports 26, 3203-3211.e5 (2019).

63. C. Hajal, et al., The CCL2-CCR2 astrocyte-cancer cell axis in tumor extravasation at the brain. Sci Adv 7 (2021).

64. F. Winkler, et al., Imaging glioma cell invasion in vivo reveals mechanisms of dissemination and peritumoral angiogenesis. Glia 57, 1306–1315 (2009).

65. T. L. Moore, et al., Nanoparticle administration method in cell culture alters particle-cell interaction. Sci Rep 9, 900 (1).

66. L. Shi, M. Zeng, Y. Sun, B. M. Fu, Quantification of blood-brain barrier solute permeability and brain transport by multiphoton microscopy. J Biomech Eng 136, 031005 (2014).

67. M. E. Berens, A. Giese, “…those left behind.” Biology and oncology of invasive glioma cells. Neoplasia 1, 208–19 (1999).

68. S. V. Sheleg, et al., Local chemotherapy with cisplatin-depot for glioblastoma multiforme. J Neurooncol 60, 53–9 (2002).

69. M. B. Chen, et al., Inflamed neutrophils sequestered at entrapped tumor cells via chemotactic confinement promote tumor cell extravasation. Proc Natl Acad Sci U S A 115, 7022–7027 (7).

70. K. Haase, M. R. Gillrie, C. Hajal, R. D. Kamm, Pericytes Contribute to Dysfunction in a Human 3D Model of Placental Microvasculature through VEGF-Ang-Tie2 Signaling. Adv Sci (Weinh) 6, 1900878 (2019).

71. J. Schindelin, et al., Fiji: an open-source platform for biological-image analysis. Nat Methods 9, 676–82 (2012).

72. P. Bankhead, et al., QuPath: Open source software for digital pathology image analysis. Sci Rep 7, 16878 (12).

73. G. S. Offeddu, et al., The cancer glycocalyx mediates intravascular adhesion and extravasation during metastatic dissemination. Commun Biol 4, 255 (2021).

74. G. Offeddu, et al., Glycocalyx-Mediated Vascular Dissemination of Circulating Tumor Cells. bioRxiv, 2020.04.28.066746 (2020).

75. S. Correa, N. Boehnke, E. Deiss-Yehiely, P. T. Hammond, Solution Conditions Tune and Optimize Loading of Therapeutic Polyelectrolytes into Layer-by-Layer Functionalized Liposomes. ACS Nano 13, 5623–5634 (2019).

76. S. Correa, et al., Highly scalable, closed-loop synthesis of drug-loaded, layer-by-layer nanoparticles. Adv Funct Mater 26, 991–1003 (2016).

77. N. Boehnke, K. J. Dolph, V. M. Juarez, J. M. Lanoha, P. T. Hammond, Electrostatic Conjugation of Nanoparticle Surfaces with Functional Peptide Motifs. Bioconjug Chem 31, 2211–2219 (9).

78. B. L. Carlson, J. L. Pokorny, M. A. Schroeder, J. N. Sarkaria, Establishment, maintenance and in vitro and in vivo applications of primary human glioblastoma multiforme (GBM) xenograft models for translational biology studies and drug discovery. Curr Protoc Pharmacol Chapter 14, Unit 14.16 (2011).

79. H. J. Kim, W. Kim, Method of tumor volume evaluation using magnetic resonance imaging for outcome prediction in cervical cancer treated with concurrent chemotherapy and radiotherapy. Radiat Oncol J 30, 70–7 (2012).

