## Supplemental information and figures for "A predictive microfluidic model of human glioblastoma to assess trafficking of blood-brain barrier penetrant nanoparticles"

### Supplementary Materials and Methods

**Cell culture and treatments.** Human iPS-ECs (Fujifilm Cellular Dynamics, 11713), human brain PCs (Catalog #1200, ScienCell), and human brain ACs (Catalog #1800, ScienCell) were cultured as described previously *(1, 2)* and used at passage 5 for all experiments. High-grade glioma patient-derived xenograft (PDX) line glioblastoma 22 (GBM22, Sarkaria lab, Mayo Clinic) was maintained in Dulbecco's Modified Eagle's Medium (DMEM) with 4.5 g/L glucose+L-glutamine without sodium pyruvate (Corning 10-017-CV) supplemented with 10% Fetal Bovine Serum (FBS) (Catalog # 26140-079, Invitrogen). For imaging, GBM22 were stably transduced with LentiBrite™ GFP Control Lentiviral Biosensor (Millipore Sigma) (gift from the Yaffe and Hammond labs, MIT), and cell line identity was confirmed by short tandem repeat (STR) testing.

**Tumor spheroid formation.** GBM22 and brain PCs, cultured in 2D, were each resuspended at 100,000 cells per mL of DMEM (Catalog # 11995073, Gibco) supplemented with 10% FBS (Catalog # 26140-079, Invitrogen). Cells were cultured in a low-adhesion 96-well plate (PrimeSurface 96M plate, Sbio) with an initial density of 5,000 cells per spheroid. For each spheroid, GBM22 and PCs were mixed at a ratio of 4:5 and a 50 µL-suspension was used per well. PCs were mixed with GBM cells in the spheroids to recapitulate tumor-stromal cell ratios commonly observed in solid tumors *(3)* and ensure the spheroids remain compact in the low-adhesion plates. DMEM (Catalog # 11995073, Gibco) with 10% FBS (Catalog # 26140-079, Invitrogen) was added to each well before centrifuging for 2 minutes at 1,000 RPM. Spheroids formed over several days by self-aggregation and were cultured for 6-7 days prior to seeding in the microfluidic devices as previously described *(1, 2)*.

**Device fabrication and microvascular network (MVN) formation.** The 3-dimensional microfluidic devices employed in this study were fabricated using soft lithography as previously described *(4, 5)* with dimensions outlined in detail elsewhere *(6)*. Briefly, a larger device with a width of 3 mm for the central gel channel and height of 500 µm was employed to ensure that spheroids can occupy the center of the channel with sufficient space for the formation of surrounding MVNs. To recapitulate the *in vivo* organization of glioma tumors surrounded by brain capillaries, spheroids were carefully removed from the 96-well plate and mixed with iPS-ECs, PCs, and ACs in fibrinogen at the ratios needed to generate the tri-culture BBB MVNs. An equal amount of thrombin was added and mixed with all the cells and spheroid prior to injecting into the devices for fibrin polymerization *(5)*. Each device was checked to ensure that the spheroid was in the center of the gel channel, with enough space for MVNs to develop and surround the tumor. Control devices were seeded with iPS-ECs, PCs, and ACs in a fibrinogen-thrombin gel without tumor spheroids. Devices were cultured in VascuLife media (Lifeline Cell Technology) supplemented with 10% FBS and 2% L-glutamine . For the first four days of culture, this media was supplemented with 50 ng/mL of VEGF-A (Catalog #100-20, Peprotech) to promote vasculogenesis following cell seeding.

**Tumor growth, vessel coverage, and vessel density measurements.** Tumor growth in the devices was measured daily between days 0 (seeding) and 7. 2-D collapsed images of the GFP signal from GBMs were acquired using an Eclipse Ti episcope (Nikon). The collapsed area of the spheroids was measured using the Fiji distribution of ImageJ (NIH) *(7)* and quantified over the course of culture. Vessel density was computed as previously described *(8)*. Using confocal microscopy (FV-1200, Olympus), 3-dimensional images of GFP-labeled GBM spheroids surrounded by BBB vessels with iPS-ECs stained for CD31 (ab3245, Abcam) were obtained. Concentric regions of interest were generated at 150 µm (proximal) and 300 µm (distal) from the surface of the tumor spheroid. Vessels were projected in the z-direction and converted to binary images to calculate the respective percent area coverage of vessels in each region of interest: tumor center, proximal, and distal ROI. These results were compared to prior results from our group using human umbilical vein endothelial cell (HUVEC) MVNs and ovarian (Skov3) spheroids or A549 (lung) spheroids *(8)*.

**Immunostaining details.** Devices were fixed with 4% PFA for 15 minutes and permeabilized with 0.01% Triton X-100 for 5 minutes. Devices were then blocked with 4% w/v BSA (Catalog # A9647, Millipore Sigma) and 0.5% v/v goat serum (Catalog # 16210064, Gibco) in PBS overnight at 4C on a shaker. Following a PBS wash, protein visualization was achieved by staining the fixed devices with anti-CD31 (ab3245, Abcam), anti-LRP1 (sc-57351, Santa Cruz Biotechnology), and anti-ZO-1 (61-7300, ThermoFisher Scientific) at 1:200 in PBS, overnight at 4C on a shaker. Following another PBS wash, devices were incubated with secondary antibodies at 1:200 in PBS (568 goat anti-rabbit A-11011, or 633 goat anti-mouse A-21052, Invitrogen) and DAPI (D1306, Invitrogen) at 1:1000 in PBS, overnight at 4C on a shaker. Images were acquired with a confocal laser scanning microscope (FV-1200, Olympus).

For in vivo samples, brains were formalin fixed and paraffin embedded before sectioning into 5 µm thick samples. Cleaved caspase-3 staining (CC3 Rabbit Mab, 1:800 #9664L [D175] Cell Signaling Technology) was performed at pH6 in heat induced epitope retrieval citrate buffer, 60 minutes at room temperature. After washing, rabbit polymer secondary (Biocare Medical # RMR 622L) was incubated for 30 minutes followed by 3,3'-diaminobenzidine for 5 minutes.

**Nanoparticle synthesis, layering, and cisplatin quantification.** Fluorescent liposomes were generated using thin film hydration followed by extrusion as previously described *(9)*. Cholesterol, 1,2-distearoyl-sn-glycero-3-phosphocholine (DSPC), 1,2-distearoyl-sn-glycero-3-phosphoethanolamine (DSPE) and 1,2-distearoyl-sn-glycero-3-phospho-(1'-rac-glycerol) (DSPG, all lipids from Avanti Polar Lipids, Alabaster, AL) were dissolved in organic solvents at 31 DSPC, 31 cholesterol, 31 DSPG and 6 DSPE (all mol%). After rehydration, liposomes were extruded using an Avestin LiposoFast LF-50 liposome extruder through sequentially smaller nucleopore membranes until liposomes with diameter 50-100nm were achieved.

For cisplatin-loaded liposomes, the film was rehydrated with a highly concentrated 8mg/ml solution of cisplatin (CDDP) in milliQ water at 80°C prior to extrusion. Next, liposomes were fluorescently labeled through NHS-coupling of sulfo-cyanine5 NHS ester dye (Lumiprobe, Hunt Valley MD) to DSPE headgroups according to the manufacturer instructions. Excess dye and/or drug was removed via KrosFlo II tangential flow filtration (TFF) system (Repligen, Waltham, MA). Cisplatin was quantified using an Agilent 1200 Series Gradient high performance liquid chromatography (HPLC) System (Agilent, Santa Clara CA) after derivatization with sodium diethyldithiocarbamate trihydrate (DTTC, Sigma Aldrich). Briefly, cisplatin-containing nanoparticles were dissolved in methanol, complexed with DTTC, extracted into chloroform by vortexing and centrifuging. After evaporating organic solvent, samples were resuspended in dimethylformamide for HPLC analysis using a fixed gradient of 80/20 methanol/water and detected at 254nm wavelength. For release assays, Float-A-Lyzer dialysis devices (Repligen, Waltham, MA) were used according to manufacturer instructions in serum-containing media at 37°C or milliQ water at 4°C (storage conditions). Releasate was sampled at predetermined timepoints and cisplatin was quantified with HPLC.

Fluorescent polystyrene cores were purchased as FluoSpheres™ Carboxylate-Modified Microspheres, red fluorescent (580/605, Invitrogen) with diameters of 0.02 µm, 0.1 µm, and 0.5 µm. All NP cores had negative surface potential for electrostatic layering. Layering was achieved by sequentially adding oppositely charged polyelectrolytes as previously described *(10)*. For liposomes, layering was performed in a solution with 25mM HEPES and 20mM HCl; for polystyrene cores, layering was performed in water. Poly-(L-arginine) (PLR) was used for the cationic layer and propargyl-modified poly-(L-aspartic acid) (PLD) was used for terminal anionic layer.

**Cisplatin (CDDP) NP treatment *in vitro* and cell death quantification.** CDDP NPs were made fresh and cisplatin concentrations in each NP formulation were quantified prior to each use. BBB-GBM devices were treated daily with appropriate volumes of free CDDP or CDDP NPs to achieve a treatment concentration of 6 µM per device per day. Tumor size was measured over time with an Eclipse Ti episcope (Nikon) as previously described. At times, fluorescently-labeled CDDP NPs with cyanine5 were used to quantify CDDP NP uptake in GBM tumors over time. To evaluate cell death, devices were incubated with 5 µM of Sytox Orange (Catalog # S11368, ThermoFisher Scientific) for 1 h and DAPI, applying a hydrostatic pressure drop across the gel channel *(5)*. Devices were imaged using a confocal microscope (FV-1200). Following automatic thresholding and segmentation using the Fiji distribution of ImageJ, z-stack images were employed to generate a 3-dimensional mask of the GBM tumors to quantify Sytox signal inside and near the tumor(11)(11).

**Real-time quantitative reverse transcription polymerase chain reaction (qRT-PCR).** Gene expression was quantified via real-time qRT-PCR for BBB-GBM devices treated with free CDDP, bare CDDP NPs, or AP2 CDDP NPs. Similar to the western blot assay, the PDMS of the devices was separated from the glass coverslip using a razor blade, and the GBM tumor region was collected as well as a region far away from the GBM spheroid for each device. 3 devices were pooled for each condition into one sample and n = 3 samples were used for each condition (n = 9 devices total). The collected samples were immersed in 50 µL of liberase resuspension medium maintained on ice for 30 minutes. Liberase resuspension medium was prepared by dissolving liberase (Catalog # 5401119001, Millipore Sigma) in sterile water to a final concentration of 5 mg/mL before storing at -80 °C. Total RNA was isolated and purified using the RNeasy Mini Kit (74104, Qiagen) according to the manufacturer's instructions. The concentration of total RNA was measured using a NanoDrop 1000 spectrophotometer. cDNA was synthesized using the High-Capacity RNA-to-cDNA Kit (4387406, Thermo Fisher Scientific) according to the manufacturer's protocol. Real-time qRT-PCR was performed on the 7900HT Fast Real-Time PCR System using the TaqMan Fast Advanced Master Mix (4444556, Thermo Fisher Scientific) as previously described *(2)*. Glyceraldehyde phosphate dehydrogenase (GAPDH) was used as a housekeeping gene, and the following genes with respective TaqMan Gene Expression Assays (Thermo Fisher Scientific) were used: Annexin V (Hs00996186_m1), Caspase 3 (Hs00234387_m1), and Caspase 7 (Hs00169152_m1).

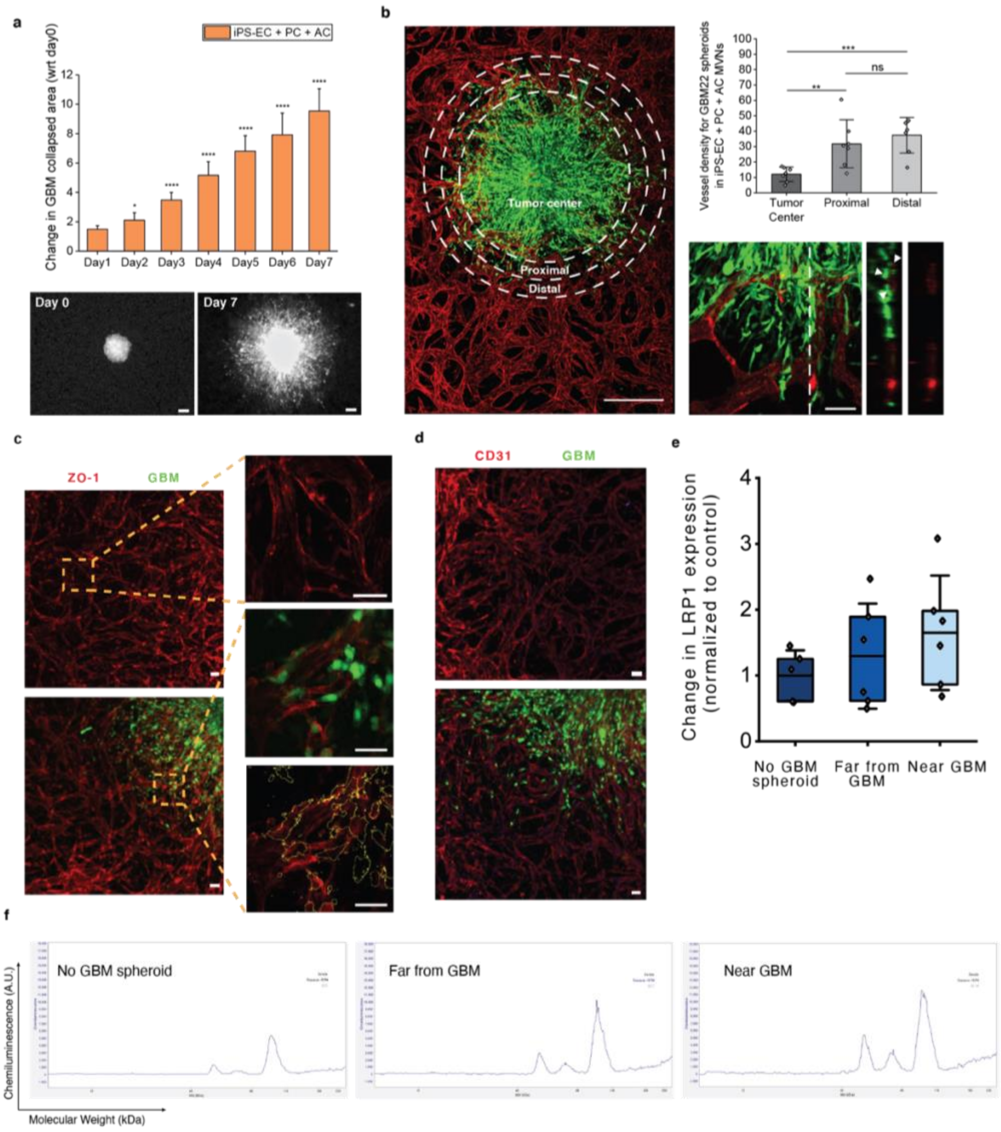


**Fig. S1. BBB microvascular network (MVN) platform characterization with GBM tumor spheroids. (A)** GBM tumor size increase over 7 days of culture in the microfluidic devices with BBB MVNs. Each datapoint represents 1 GBM tumor grown in 1 device with BBB MVNs, scale bar = 100µm. **(B)** Representative images of a GBM tumor grown in BBB MVNs and its ability to co-opt the vasculature. Vessel density is quantified in 3 regions of interest: tumor center (> 80 % tumor surface coverage), proximal area (300µm away from tumor center area), and distal area (300µm away from proximal area). Each datapoint represents 1 device. **(C)** Validation of tight junction (ZO-1) protein expression in iPS-ECs for control BBB MVNs (no GBM, top panel) and BBB MVNs with GBM tumors (bottom panel, the same zoomed region is shown with GBM signal (GFP) and without GFP signal but with yellow contour for GBM cells). ZO-1 expression is not altered near GBM tumors, particularly in vessel branches were GBM co-option is observed. **(D)** Validation of adherens junction (CD31) protein expression in iPS-ECs for control BBB MVNs (no GBM) and BBB MVNs with GBM tumors. **(E)** Change in LRP1 protein expression in three regions of interest within the BBB-GBM device, plotted normalized to *No GBM spheroid* control; western blot quantitative data was normalized to CD31 for each sample to control; data points represent n=1 device. **(F)** Example of raw, uncropped chemiluminescence signals for LRP1 expression in n=1 device, as quantified in e for n=6 devices.

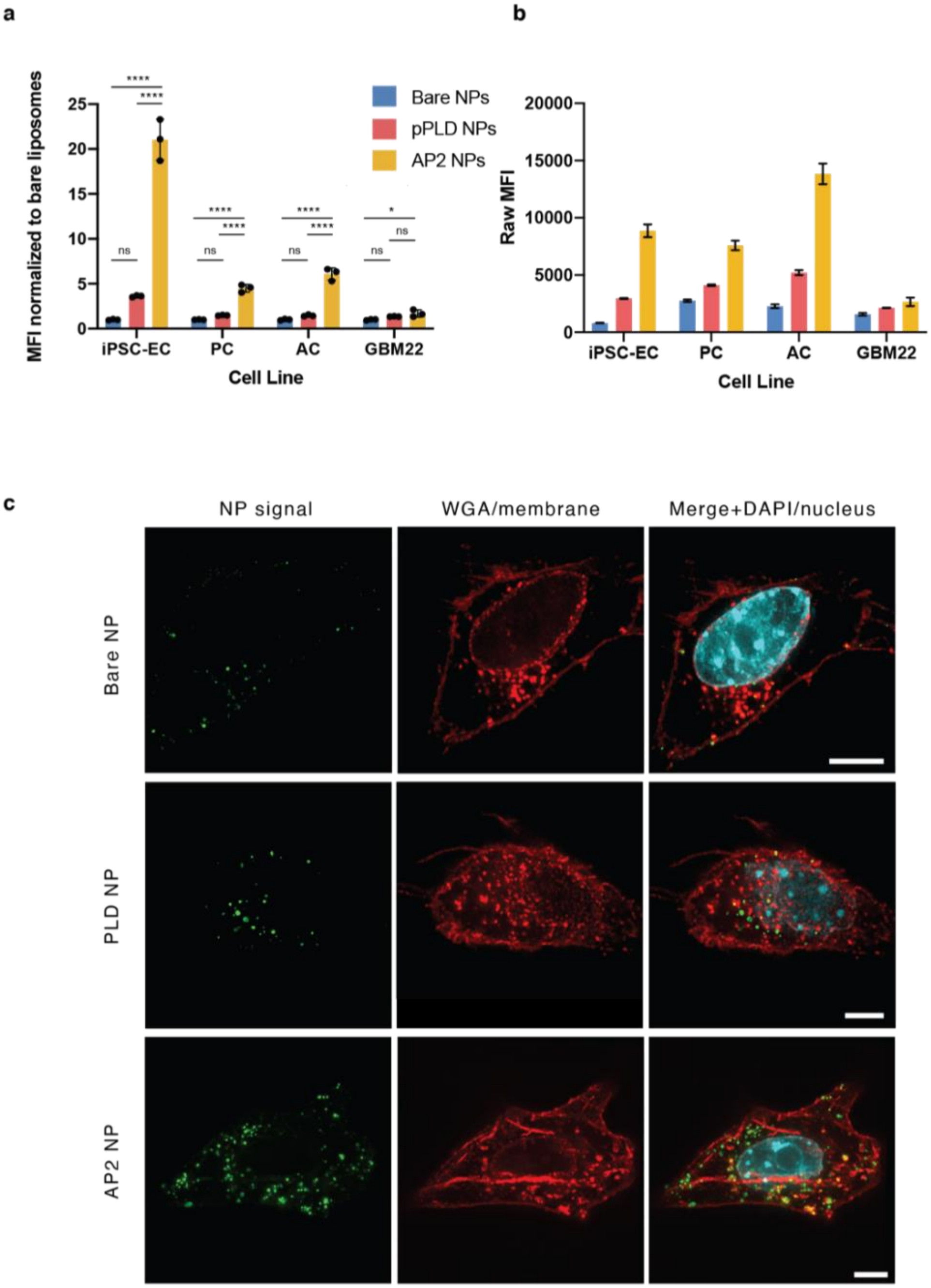


**Fig. S2. Differential nanoparticle (NP)-cell association of functionalized NPs in blood-brain barrier component cells and GBM cells *in vitro*.** (**A**) NP-cell association measured using flow cytometry after 24hr incubation with induced pluripotent stem cell endothelial cells (iPSC-ECs), pericytes (PCs), astrocytes (ACs), and GBM22 glioblastoma cell line, normalized to Bare NP control within each cell type. NPs tested as bare liposomes (Bare NPs), layered with propargyl poly-(L-aspartic acid) (pPLD NPs) and angiopep-2 conjugated NPs (AP2-NPs). Bars represent mean ± standard deviation (S.D.), n.s. denotes not significant, * $p < 0.05$, ** $p < 0.01$, *** $p < 0.001$, and **** $p < 0.0001$ using ANOVA with Sidak correction for multiple comparisons. (**B**) Raw median fluorescence intensity (MFI) for data presented in a. (**C**) Representative micrographs of NP formulations in GBM22 cells after 24hr incubation which NP signal is pseudo colored green, cell nucleus in cyan and cell membrane in red, highlighting internalization of NPs after surface functionalization. Scale bars = 5 µm.

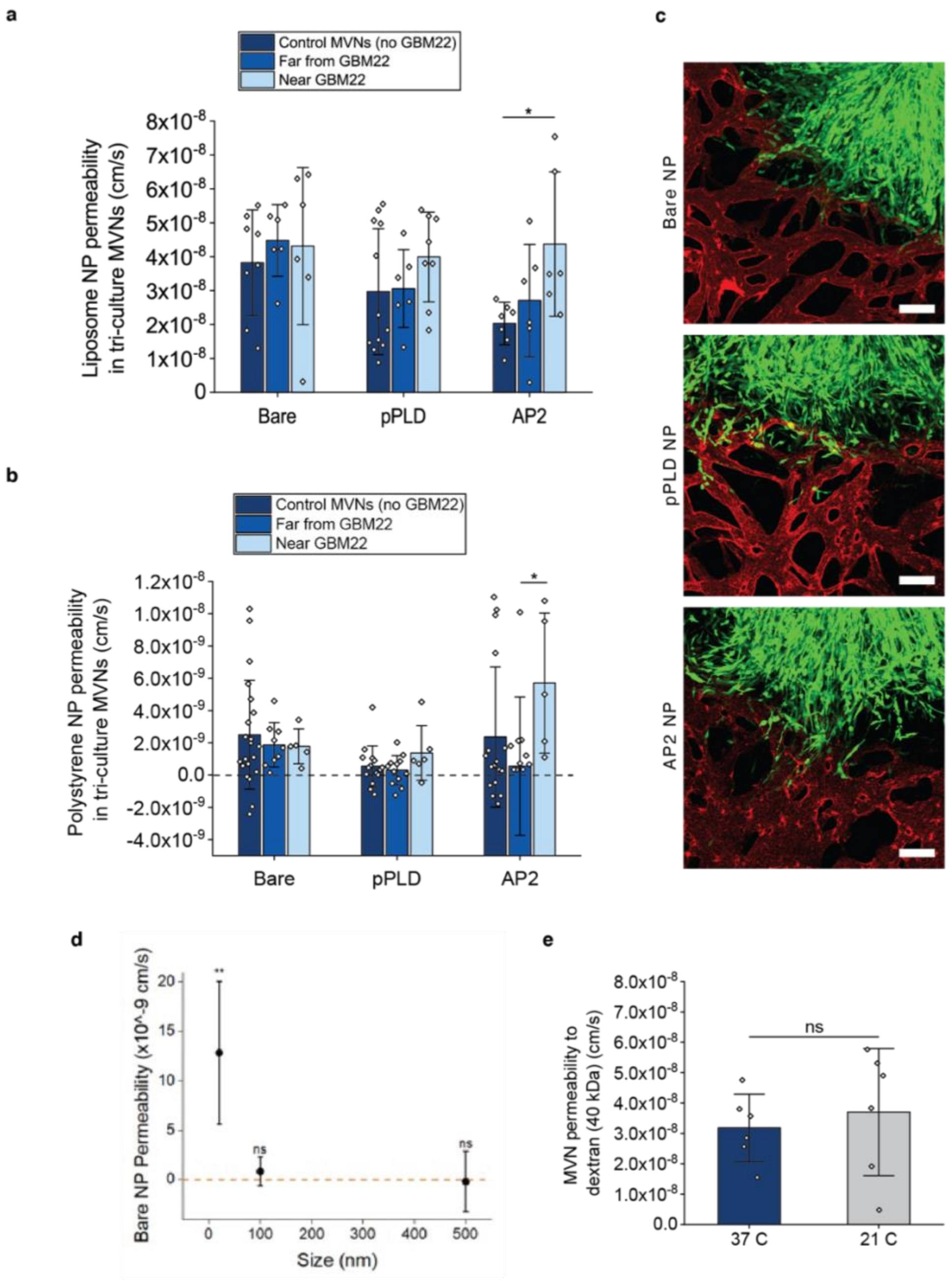


**Fig. S3. NP permeability measurements in the BBB microvascular networks (MVNs) with or without GBM tumors for liposomal and polystyrene NPs.** (**A**) Absolute values of permeability for liposomal NPs presented in main Figure 2D. (**B**) Polystyrene NP permeability measurements in BBB MVNs in the 3 regions of interest (no GBM, far from GBM, and near GBM) for the 3 NP formulations considered. 100nm polystyrene NPs were employed to generate bare, pPLD, and AP2 formulations. Each datapoint represent n = 1 ROI; n = 6 independent devices per condition were considered. (**C**) Representative images of polystyrene NPs in the BBB microvessels near GBM spheroid at t = 0 min following NP perfusion; time-lapse images over 12 min were used to determine permeabilities in (C); GBM spheroids are psuedocolored green and nanoparticles are in red; scale bars =100 µm. (**D**) Permeability measurements as a function of NP size in BBB MVNs without GBM tumors. Bare polystyrene NPs of 20, 100, and 500nm diameters were used to obtain permeability measurements. N = 3 devices per NP size. (**E**) Permeability to 40 kDa dextran in the BBB MVNs (no GBM tumor) at 37 or 21 °C.

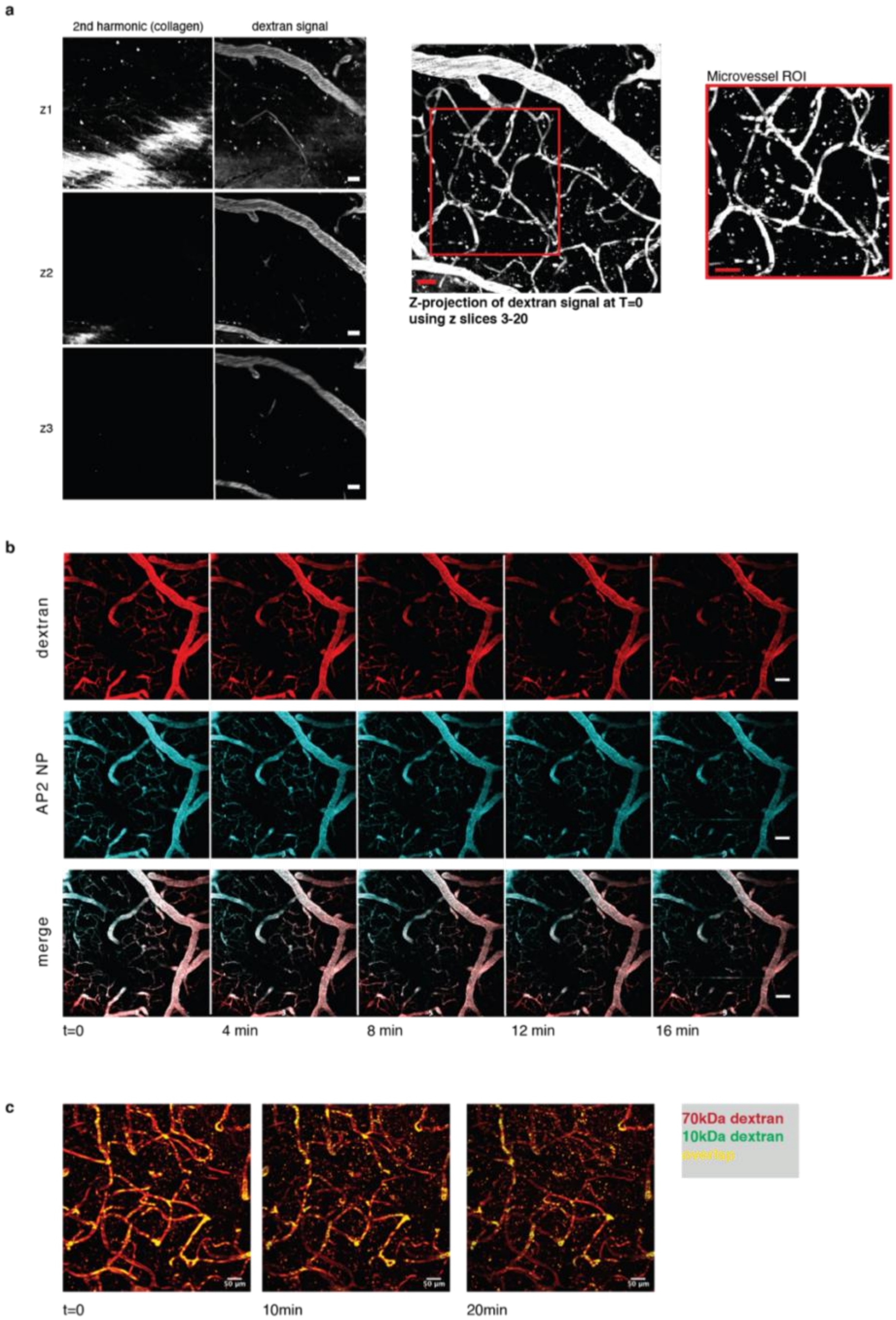


**Fig. S4. Technical details of intravital imaging technique** (**A**) Images were acquired such that the top z-slice contained second harmonic signal representing thinned bone. Z-slices were 10µm apart and the top 1-2 z-slices (containing collagen) were excluded prior to analysis to ensure that no scalp vessels were included. Choice of region of interest (ROI) to evaluate capillary-sized micro vessels for permeability measurements was made using a z-projection (excluding top z-slices); the red outlined inset shows a representative microvessel ROI at time = 0. (**B**) Example of time-lapse z-projections from t=0 to t=16 minutes for 40kDa dextran (red) and AP2 nanoparticles (cyan). Images were acquired in a 100µM z-stack at 2 minute intervals. (**C**) Time-lapse z-projection images after co-administration of 70 kDa and 10 kDa dextran highlights differential dwell time within the vessels, with decreasing areas of overlap and increased 10 kDa dextran signal in the brain parenchyma during the 20 minute imaging period. Throughout, scale bars = 50µm.

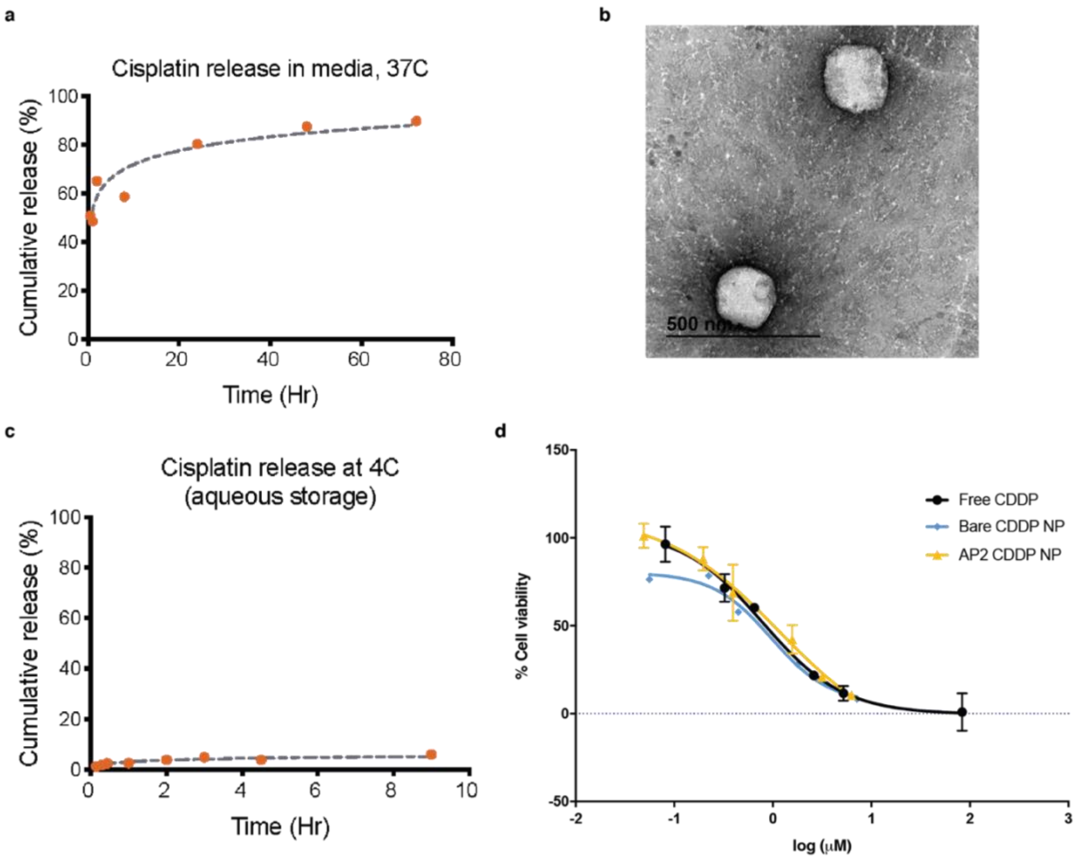


**Fig. S5. Characterization of cisplatin-loaded nanoparticles.** (**A**) Cumulative release of cisplatin (CDDP) from drug-loaded layer-by-layer nanoparticles at 37°C in cell media. (**B**) Representative transmission electron micrograph of CDDP-loaded liposome with poly-(L-arginine) layer; scale bar = 500nm. (**C**) Cumulative release of cisplatin (CDDP) from drug-loaded layer-by-layer nanoparticles at 4C in water (storage conditions). (**D**) Drug-response curves for GBM22 cells after treatment with free CDDP, bare CDDP nanoparticles, or layer-by-layer CDDP nanoparticles with angiopep-2 (AP2) surface conjugation showing similar effect with free or encapsulated drug.

**Table S1. Nanomaterials characterization** for all functionalized and/or drug-loaded nanoparticles used in the study including z-average intensity-weighted hydrodynamic size (in nm), number-weighted average diameter (in nm), polydispersity index (PDI), nanoparticle surface zeta potential (in mV), n=3 per condition. For applicable nanoparticles, cisplatin (CDDP) drug loading is also shown (n= 3 experiments).

| **Nanoparticle Condition** | **Z-Average (d.nm)** | | **Number Mean (d.nm)** | | **Polydispersity Index (PDI)** | | **ZP (mV)** | | **Weight loading (mg CDDP/mg lipid)** | | **CDDP concentration in 1mg/ml NP solution (μM)** | |
|---|---|---|---|---|---|---|---|---|---|---|---|---|
| | AVG | SD | AVG | SD | AVG | SD | AVG | SD | AVG | SD | AVG | SD |
| **Liposomal NPs (non-drug loaded)** | | | | | | | | | | | | |
| Post extrusion (50nm) and Cy5 labeling | 106.20 | 3.00 | 68.70 | 8.90 | 0.14 | 0.04 | -45.00 | 6.27 | | | | |
| PLR layer pre-purification | 146.10 | 3.10 | 99.10 | 4.10 | 0.15 | 0.03 | 51.10 | 2.30 | | | | |
| PLR layer post-purification | 159.80 | 1.30 | 100.20 | 9.40 | 0.16 | 0.03 | 50.80 | 2.10 | | | | |
| propargyl PLD layer pre-purification | 158.20 | 2.10 | 98.90 | 2.50 | 0.19 | 0.00 | -57.80 | 3.80 | | | | |
| propargyl PLD layer post-purification | 164.80 | 3.10 | 101.40 | 12.60 | 0.16 | 0.02 | -45.40 | 0.80 | | | | |
| AP2-clicked (post purification) | 163.50 | 2.60 | 95.90 | 22.40 | 0.16 | 0.03 | -36.50 | 1.00 | | | | |
| **Polystyrene NPs (non-drug loaded)** | | | | | | | | | | | | |
| Received from manufacturer | 117.20 | 0.75 | 95.34 | 1.85 | 0.02 | 0.01 | -42.67 | 0.40 | | | | |
| PLR layer pre-purification | 125.67 | 0.99 | 95.00 | 6.95 | 0.06 | 0.04 | 47.50 | 1.22 | | | | |
| PLR layer post-purification | 167.57 | 3.03 | 111.17 | 19.25 | 0.11 | 0.07 | 75.00 | 0.82 | | | | |
| propargyl PLD layer pre-purification | 145.70 | 3.86 | 99.95 | 10.42 | 0.10 | 0.05 | -48.97 | 0.55 | | | | |
| propargyl PLD layer post-purification | 148.07 | 2.62 | 112.60 | 8.71 | 0.08 | 0.04 | -58.53 | 1.03 | | | | |
| AP2-clicked (post purification) | 146.33 | 1.83 | 104.11 | 11.08 | 0.16 | 0.03 | -36.50 | 0.82 | | | | |
| **Cisplatin-loaded liposomal NPs** | | | | | | | | | | | | |
| Post extrusion (50nm) and drug loading | 104.13 | 1.04 | 71.22 | 11.28 | 0.18 | 0.02 | -46.40 | 0.85 | 6.03 | 1.72 | 201.11 | 57.57 |
| PLR layer pre-purification | 113.87 | 2.22 | 79.93 | 1.51 | 0.14 | 0.01 | 45.37 | 1.58 | | | | |
| PLR layer post-purification | 117.37 | 1.65 | 78.34 | 6.62 | 0.16 | 0.04 | 54.27 | 0.60 | | | | |
| propargyl PLD layer pre-purification | 122.27 | 1.58 | 87.65 | 2.54 | 0.15 | 0.05 | -56.37 | 1.92 | | | | |
| propargyl PLD layer post-purification | 125.73 | 1.33 | 96.07 | 4.21 | 0.13 | 0.02 | -49.23 | 0.87 | | | | |
| AP2-clicked (post purification) | 125.70 | 2.76 | 85.77 | 5.09 | 0.18 | 0.06 | -57.77 | 1.97 | 3.50 | 1.26 | 116.56 | 42.02 |

**Table S2. Morphologic parameters** of the *in vitro* BBB model and *in vivo* mouse BBB capillaries described in this paper.

| | 3D in vitro BBB model (iPS-EC + PC + AC)<br>(n=6 devices) | In vivo mouse BBB capillaries<br>(n = 6 mice) |
|---|---|---|
| Average diameter (um) | (33.54 ± 1.06) | (3.76 ± 0.65) |
| % vessel coverage | (7.77 ± 3.90) | (11.69 ± 3.69) |
| Branch length (um) | (174.28 ± 13.70) | (8.97 ± 1.69) |
| # of branches per area | (39.35 ± 5.83) | (418.82 ± 194.23) |

**Table S3.** Comparison of *in vitro* and *in vivo* dextran permeability values from this paper to *in vivo* permeabilities in similar published works.

| Dextran Molecular Weight | 3D in vitro BBB device (iPS-EC + PC + AC)<br><br>Source: this paper | In vivo mouse BBB capillaries<br><br>Source: this paper | In vivo rat BBB capillaries (multiphoton)<br><br>Source: Yuan et al., 2009 (12) | In vivo rat BBB capillaries (inverted microscope)<br><br>Source: Kutuzov et al., 2018 (13) |
|---|---|---|---|---|
| 40 kDa | $(3.18 \pm 1.11) \times 10^{-8}$ cm/s | $(4.97 \pm 2.37) \times 10^{-8}$ cm/s | $(1.37 \pm 0.26) \times 10^{-7}$ cm/s | $(1.9 \pm 1.1) \times 10^{-7}$ cm/s |
| 70 kDa | Not quantified | $(1.52 \pm 1.96) \times 10^{-8}$ cm/s | $(1.30 \pm 0.20) \times 10^{-7}$ cm/s | $(1.5 \pm 0.5) \times 10^{-7}$ cm/s |

**Movie S1 (separate file).** Two-photon microscopy time-lapse imaging of 40kDa dextran (left panel), bare Cy5-labeled nanoparticle signal (center panel), and merged images (right panel), all shown as maximum z-projections of 10 images over total depth of 100µm, images taken over 20 minutes at 2 minute intervals. Scale bar =50µm.

**SI References**


1. M. Campisi, *et al.*, 3D self-organized microvascular model of the human blood-brain barrier with endothelial cells, pericytes and astrocytes. *Biomaterials* **180**, 117–129 (10).
2. C. Hajal, *et al.*, The CCL2-CCR2 astrocyte-cancer cell axis in tumor extravasation at the brain. *Sci Adv* **7** (2021).
3. S. V. Sheleg, *et al.*, Local chemotherapy with cisplatin-depot for glioblastoma multiforme. *J Neurooncol* **60**, 53–9 (2002).
4. F. Winkler, *et al.*, Imaging glioma cell invasion in vivo reveals mechanisms of dissemination and peritumoral angiogenesis. *Glia* **57**, 1306–1315 (2009).
5. M. B. Chen, *et al.*, Inflamed neutrophils sequestered at entrapped tumor cells via chemotactic confinement promote tumor cell extravasation. *Proc Natl Acad Sci U S A* **115**, 7022–7027 (7).
6. K. Haase, M. R. Gillrie, C. Hajal, R. D. Kamm, Pericytes Contribute to Dysfunction in a Human 3D Model of Placental Microvasculature through VEGF-Ang-Tie2 Signaling. *Adv Sci (Weinh)* **6**, 1900878 (2019).
7. J. Schindelin, *et al.*, Fiji: an open-source platform for biological-image analysis. *Nat Methods* **9**, 676–82 (2012).
8. K. Haase, G. S. Offeddu, M. R. Gillrie, R. D. Kamm, Endothelial Regulation of Drug Transport in a 3D Vascularized Tumor Model. *Adv Funct Mater* **30** (2020).
9. S. Correa, N. Boehnke, E. Deiss-Yehiely, P. T. Hammond, Solution Conditions Tune and Optimize Loading of Therapeutic Polyelectrolytes into Layer-by-Layer Functionalized Liposomes. *ACS Nano* **13**, 5623–5634 (2019).
10. S. Correa, *et al.*, Highly scalable, closed-loop synthesis of drug-loaded, layer-by-layer nanoparticles. *Adv Funct Mater* **26**, 991–1003 (2016).
11. G. S. Offeddu, *et al.*, An on-chip model of protein paracellular and transcellular permeability in the microcirculation. *Biomaterials* **212**, 115–125 (2019).
12. W. Yuan, Y. Lv, M. Zeng, B. M. Fu, Non-invasive measurement of solute permeability in cerebral microvessels of the rat. *Microvasc Res* **77**, 166–73 (2009).
13. N. Kutuzov, H. Flyvbjerg, M. Lauritzen, Contributions of the glycocalyx, endothelium, and extravascular compartment to the blood-brain barrier. *Proc Natl Acad Sci U S A* **115**, E9429–E9438 (10).
